# Stem cell specific interferon stimulated gene expression is regulated by the formative pluripotency network through IRF1

**DOI:** 10.1101/2021.12.07.471598

**Authors:** Merrit Romeike, Stephanie Spach, Marie Huber, Songjie Feng, Gintautas Vainorius, Ulrich Elling, Christa Buecker

## Abstract

Stem cells intrinsically express a subset of genes which are normally associated with interferon stimulation, and thus the innate immunity response. Expression of these interferon stimulated genes (ISG) in stem cells is independent of external stimuli such as viral infection. Here we show that the interferon regulatory factor 1, *Irf1*, is directly controlled by the murine formative pluripotency gene regulatory network and therefore upregulated in the transition from naive to formative pluripotency. IRF1 in turn binds to regulatory regions of a conserved set of ISGs and is required for their faithful expression in formative pluripotent cells. IRF1 also binds to an enhancer of the formative pluripotency transcription factor *Oct6* and is partially required for upregulation of *Oct6*. IRF1 therefore acts as a link between the formative pluripotency network and the regulation of innate immunity genes in formative pluripotency.

## Introduction

Mouse embryonic stem cells (ESCs) are self-renewing in a naive pluripotent cell state *in vitro*. The essential gene regulatory network required to maintain naive pluripotency has been the subject of numerous studies and is very well defined (Betschinger et al. 2013; Dunn et al. 2014; Leeb et al. 2014). In contrast to maintenance of one cell state, development is characterized by tightly regulated cell fate transitions, which are required to establish the remarkable complexity of multicellular organisms. After exiting naive pluripotency, cells enter a transient state termed formative pluripotency (Smith 2017). This transition is characterized by global reorganization of the enhancer landscape and upregulation of the formative gene regulatory network (Buecker et al. 2014). However, factors involved in this formative pluripotency gene regulatory network are less well understood: genetic screens interrogating this transition have mostly focused on maintenance of (Li et al. 2018; Seruggia et al. 2019) or exit from naive pluripotency (Betschinger et al. 2013; Leeb et al. 2014; Li et al. 2018; Lackner et al. 2021). Formative pluripotency was only considered in the context of a multistep differentiation into primordial germ cell fate (Hackett et al. 2018).

While it is not clear which factors are involved in the formative pluripotency network, stem cells intrinsically express a subset of genes which are referred to as interferon stimulated genes (ISGs) (Wu et al. 2018). How these ISGs are regulated is unclear, as stem cells do not respond to interferon stimulation (Burke, Graham, and Lehman 1978; Harada et al. 1990; Chen et al. 2020). At the same time, stem cells are resistant against viral infection (Swartzendruber and Lehman 1975; Wolf and Goff 2009). Whether ISG and pluripotency gene expression are functionally connected is so far not known.

Here we identify the interferon response gene *Irf1* in a CRISPR-KO screen as a regulator of an enhancer that controls the expression of *Oct6,* a formative marker gene. IRF1 has a conserved role as regulator of ISGs. Interestingly, many ISGs are differentially expressed between naive and formative pluripotency and we show here that IRF1 is directly responsible for the expression of a subset of these genes. Furthermore, *Irf1* expression in formative pluripotent cells is directly regulated by the formative pluripotency gene regulatory network through a stem cell specific enhancer. Pluripotency and interferon responsive gene regulatory networks are therefore connected by *Irf1* in the transition from naive to formative pluripotency.

## Results

### Monitoring exit from naive and entry into formative pluripotency with dual fluorescent reporter

We designed a fluorescent reporter system which simultaneously monitors exit from naive and entry into formative pluripotency. Mouse embryonic stem cells are cultured under defined 2i+LIF conditions, which preserve the naive state of pluripotency. Upon withdrawal of 2i+LIF and stimulation with FGF2, the cells differentiate within 48h irreversibly from the mESC state into Epiblast like cells (EpiLC) (Hayashi et al. 2011; Buecker et al. 2014), often referred to as the formative state of pluripotency (Smith 2017). We identified two enhancer elements that are differentially active in either the naive or the formative state of pluripotency **(Figure S1A**) and coupled them to activate different fluorescent markers to follow the exit from naive pluripotency and the entry into formative pluripotency simultaneously (**Figure 1A, S1A**)): an enhancer close to the naive transcription factor *Tbx3* to control GFP expression and an enhancer close to the formative marker gene *Oct6* (also known as *Pou3f1*) to drive mCherry expression (Buecker et al. 2014). Both constructs were cloned into piggybac vectors with different selection cassettes and transfected into ESC cultured under defined 2i+LIF conditions. We selected two independent clonal cell lines which showed clear separation of ESC and EpiLC populations after 48 h of differentiation by FACS analysis (**Figure 1B, S1B**). Next, we transfected constitutively expressed *Cas9* to enable a CRISPR-KO-based screening approach. To ensure proper function of the screening set up, we tested if the generated dual reporter screening cell lines report impaired differentiation: we deleted *Tcf7l1* (also known as *Tcf3*), a transcription factor for which deletion causes a strong delay in the exit from naive pluripotency (Wray et al. 2011). As expected, *Tcf7l1* knock-out (KO) caused a shift in dual fluorescent marker expression in the EpiLC population (**Figure 1C**), with mCherry showing lower expression compared to the WT cells and GFP showing higher expression in the *Tcf7l1* KO cells 48h after initiation of differentiation. In summary, the dual fluorescent reporter enables faithful monitoring of the transition from naive into formative pluripotency.

**Figure 1.**
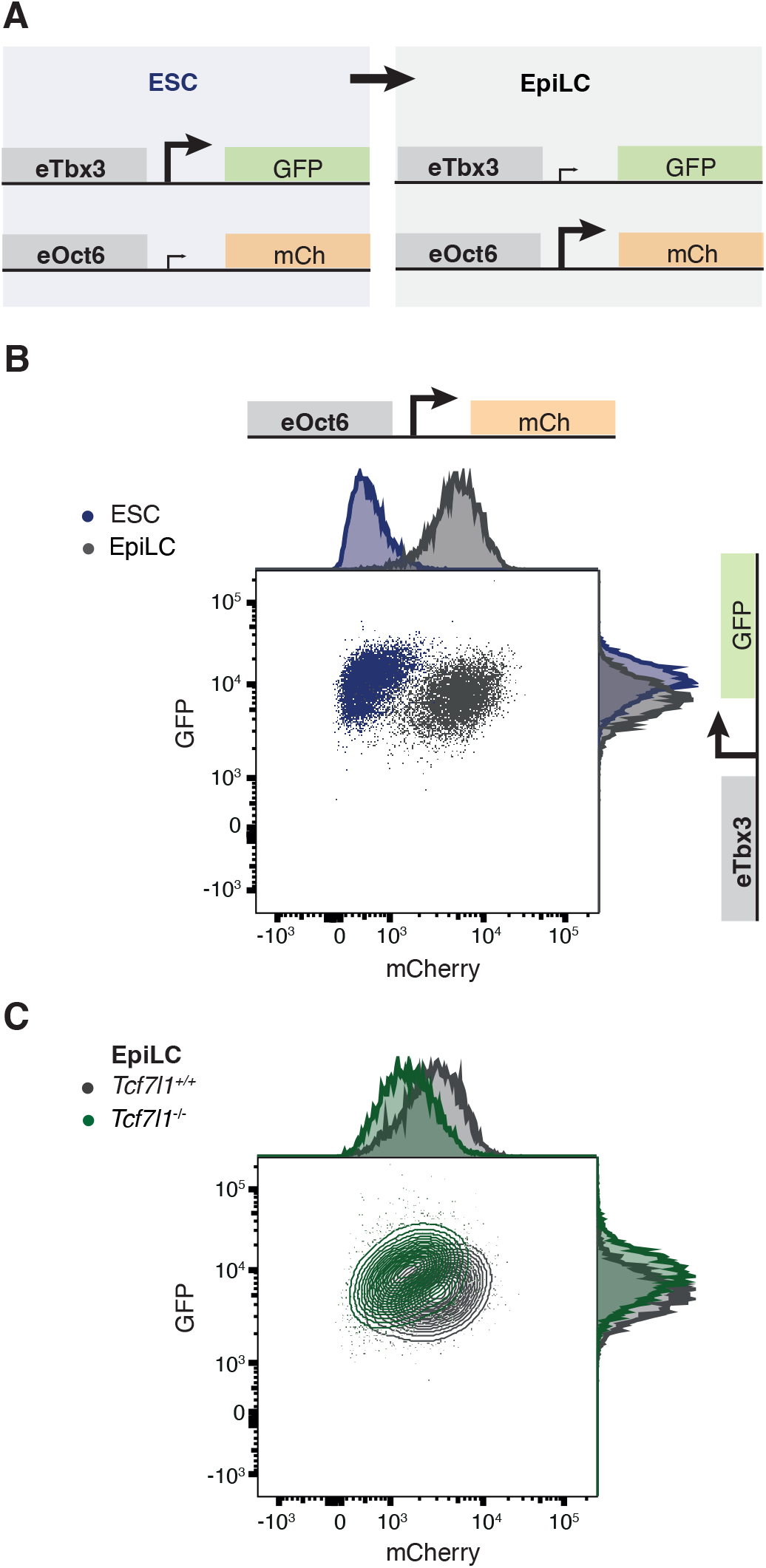
Dual fluorescent reporter cell line monitors activation of formative pluripotency. **A** Scheme of the reporter system: ESCs express GFP under *eTbx3* enhancer control and mCherry under control of the *eOct6* enhancer. When cells are differentiated to EpiLCs, GFP expression is diminished and mCherry expression increases. **B** Representative flow cytometry profiles of reporter cell lines in ESC and differentiated to EpiLC conditions. See also Figure S1B **C** Representative flow cytometry profiles of reporter cell lines in EpiLC condition, *Tcf7l1^+/+^* and *Tcf7l1*^-/-^.

### Screening for factors regulating entry into formative pluripotency

Next, we used the generated dual reporter cell line to identify factors regulating the entry into formative pluripotency by performing a pooled CRISPR-KO screen. We transduced the reporter cell lines (100 mio cells) with a lentiviral library for the expression of 22781 unique sgRNAs targeting 2524 mostly nuclear localized genes, including 159 genes considered ISGs (**Figure 2A, S2a**). We chose this smaller, selected library over a genome-wide screen to increase statistical robustness. Furthermore, we expected a candidate regulator of formative pluripotency to regulate transcription. To control for guide abundance in the lentiviral library, we also infected wildtype ESC (R1 cells) lacking CAS9 expression. We selected infected cells with Neomycin for 48 h. After recovery from selection, we harvested 60 mio cells to identify ESC essential factors: those factors which already enriched or depleted in the undifferentiated state. For this, we compared the abundance of guides in genomic DNA from CAS9 expressing or non-expressing cells. As expected, we observed enrichment of *Trp53* guides, as cells lacking P53 proliferate faster and enrich in populations of mixed clones (Sabapathy 1997) (**Figure 2B**). Conversely, ESC lacking the core pluripotency factors OCT4 (also known as POU5F1) and NANOG depleted from this population, since these factors are required for naive pluripotency (**Figure 2B**). Moreover, larger sets of known essential factors showed depletion (**Figure 2B, S2B**) (Hart et al. 2017; Li et al. 2018). We conclude that the overall screening set up was able to identify known essential regulators of the naive pluripotent stem cell state and could therefore be used to focus on differentiation.

**Figure 2.**
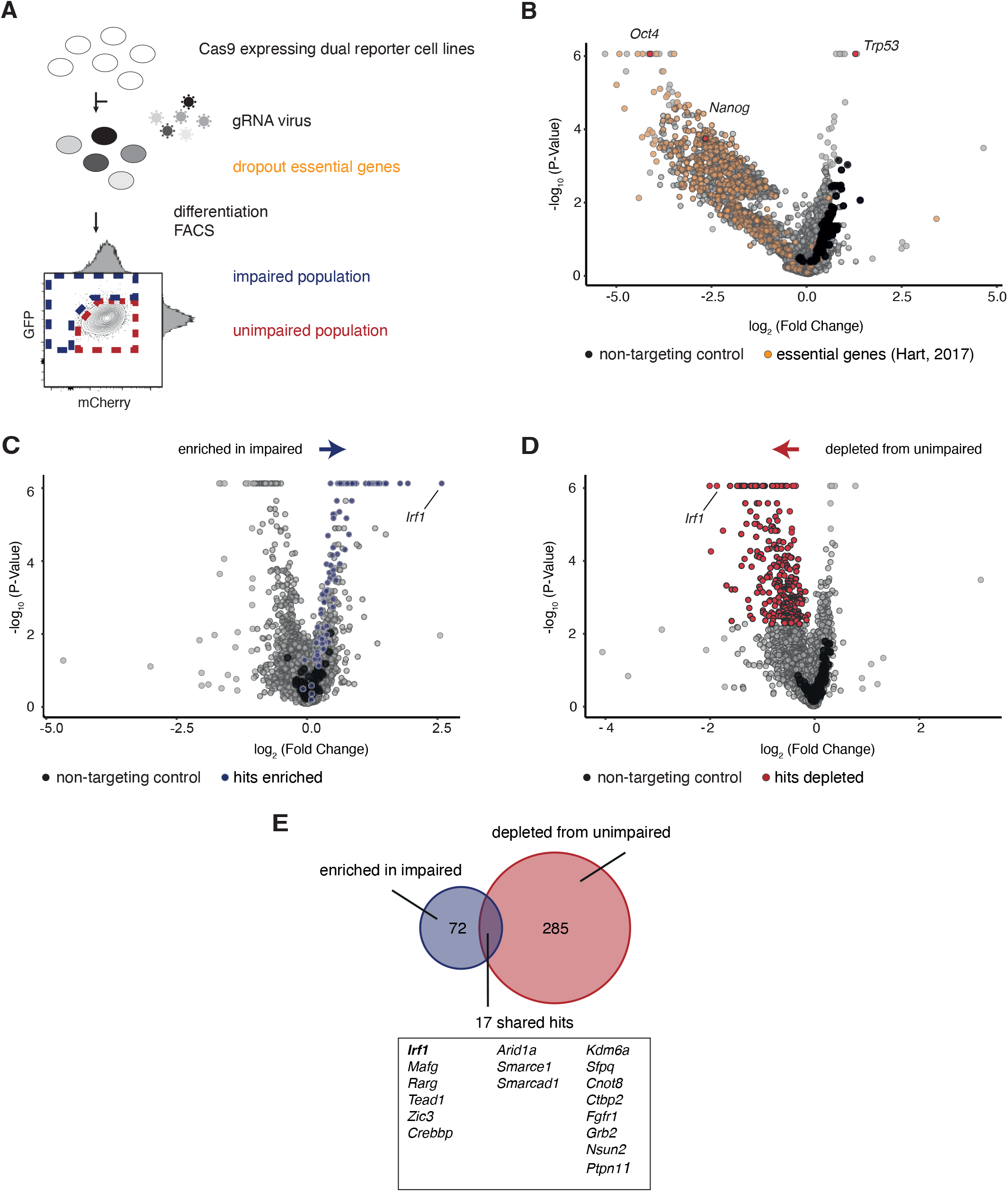
Screening for factors required for activation of formative pluripotency. **A** Scheme of pooled CRISPR-KO screening strategy. Reporter ESCs were transduced with a pooled CRISPR-KO library. Cells selected for library integration with and and without CAS9 were compared, assaying genes essential for the ESC state. Reporter cell lines were differentiated, and scored by flow cytometry sorting as differentiation impaired (blue box) or unimpaired (red box). **B-D** Volcano plots of indicated comparisons in CRISPR-KO screen analysis. The x-axis represents fold change of guide presence between indicated conditions, the y-axis shows the binomial p-value. Non-targeting control guides are indicated in black. All analyses are based on two replicates in independent cell lines. **B** Guide representation in ESCs with CAS9 vs ESCs without CAS9. Indicated are known ESC essential genes (Hart et al. 2017) and selected factors. See also Figure S2B. **C** Guide representation in EpiLC sorted as impaired vs. non-sorted EpiLC. Indicated in blue are factors enriched with FDR<0.1 **D** Guide representation in EpiLC sorted as unimpaired vs. non-sorted ESCs. Indicated in red are factors depleted with FDR<0.1 **E** Overlap of all factors enriched in impaired and/or depleted from unimpaired populations, factors found under both conditions are indicated.

We then used our screening platform to identify potential activators of the formative pluripotency network. We differentiated ESCs for 48 h into EpiLCs and binary scored EpiLCs as differentiation “impaired” or “unimpaired”: cells with reporter activity overlapping the behavior of no guide controls were FACS sorted as unimpaired. Conversely, cells with higher *eTbx3* controlled GFP and/or lower *eOct6* controlled mCherry were sorted as impaired (see **Figure 2A** for gating layout). In addition, we also collected unsorted EpiLCs as baseline. This setup allowed several comparisons of guide abundance: KO of a candidate driving the formative state should increase the probability of impaired differentiation, therefore the guides should be enriched in the impaired populations. This was the case for a total of 72 factors (**Figure 2C**). Conversely, a candidate KO should be incompatible with unimpaired differentiation and therefore guides drop out from the unimpaired population. 285 factors showed such a behavior (**Figure 2D**). We identified an overlapping set of 17 hits, which were found under both comparisons and are therefore the most stringent candidates involved in this cell fate transition (**Figure 2E**). Among these were members of pathways with known involvement in stem cell differentiation, the FGF receptor *Fgfr1* and *Tead1* which plays a role in the Hippo pathway (Molotkov et al. 2017) and *Zic3*, a known regulator of the exit from naive pluripotency and entry into primed pluripotency (Yang et al. 2019).

In addition, we identified several members of the SWI/SNF chromatin remodeling family, also called BAF complexes (*Arid1a, Smarce1, Smarcad1*). Specific subunits of BAF complexes are connected to diverse phenotypes in ESCs (reviewed in Ye, 2021). ARID1A/BAF250A and SMACRCEA/BAF57 are components of the ESC specific esBAF, with *Arid1a* deficient mice showing an arrest in early embryonic development (Gao et al. 2008). SMARCAD1 has been shown to silence endogenous retrovirus in embryonic stem cells (Sachs et al. 2019). In sum, we were able to identify known regulators of the exit from naive pluripotency showing that our screening approach can identify factors involved in the exit from naive pluripotency.

One of the top screening hits was the interferon regulatory factor, *Irf1,* a transcription factor and member of ISGs. *Irf1* was depleted from the unimpaired cell population and enriched in the impaired population for every single analyzed condition. We validated gene expression levels for all hits in ESCs and EpiLCs (**Figure S2C**). All hits are expressed, but only *Irf1* shows significant upregulation in differentiation. Upregulation of a transcription factor could indicate its function as driver of the formative pluripotency network. We therefore focused on the role of *Irf1* in the differentiation from naive to formative pluripotency.

### IRF1 KO cells fail to activate the *Oct6* enhancer

To confirm the role of IRF1 as a regulator of the activation of formative pluripotency, we generated stable *Irf1* KO cell lines.

We simultaneously transfected two CRISPR guides to delete an exon-intron boundary around intron 3 of *Irf1* into our dual reporter cell lines and R1 WT cells using lipofection. We identified individual clones through genotyping PCR and validated the absence of full-length protein by Western Blot (**Figure S3.1A**). These blots also illustrated the upregulation of *Irf1* expression in differentiation not only on RNA level (**Figure S2C**) but also on protein level (**Figure S3.1A**). Furthermore, in EpiLCs, IRF1 is localized to the nucleus, as confirmed by immunofluorescence (**Figure S3.1B**). This staining is lost upon deletion of *Irf1* (**Figure S3.1B**).

Interestingly, *Irf1* is expressed in the inner cell mass of porcine early blastocysts (Shi et al. 2020). We turned to published datasets for early embryos of mice and humans to access if expression is conserved across mammals. Cells collected at E6.5 from the mouse embryo proper mainly correspond to epiblast or primitive streak. At the same time point, cells with robust *Irf1* expressions can be found (Pijuan-Sala et al. 2019) (**Figure S3.1C**). We also tested if early embryonic *Irf1* expression is conserved in human and analysed single cell data ranging from the 8-cell stage to prior to implantation (E3-E7) (Petropoulos et al. 2016). The majority of cells analyzed for each stage were expressing *Irf1*, with decreasing levels over time (**Figure S3.1D**). Therefore, expression of *Irf1* in early embryonic development is recapitulating the *in vivo* situation and is not an artefact of the *in vitro* ESC culturing system.

We then confirmed the effect of *Irf* KO by monitoring the activity of the dual fluorescent reporter in these KO cell lines: *Irf1* KO EpiLCs shifted along the e*Oct6* enhancer controlled mCherry expression axis and showed lower fluorescent values in FACS analysis (**Figure 3A, S3.1E**). This effect was already present in ESCs, here also the basal e*Oct6* enhancer activity was reduced (**Figure S3.1E** left panel and **S3.1G** left panel). GFP levels controlled by a e*Tbx3* enhancer were not affected (**Figure 3A, S3.1E, S3.1G**). This indicates a specific effect of IRF1 presence on the establishment of formative pluripotency, but no general effect on exit from naive pluripotency.

**Figure 3.**
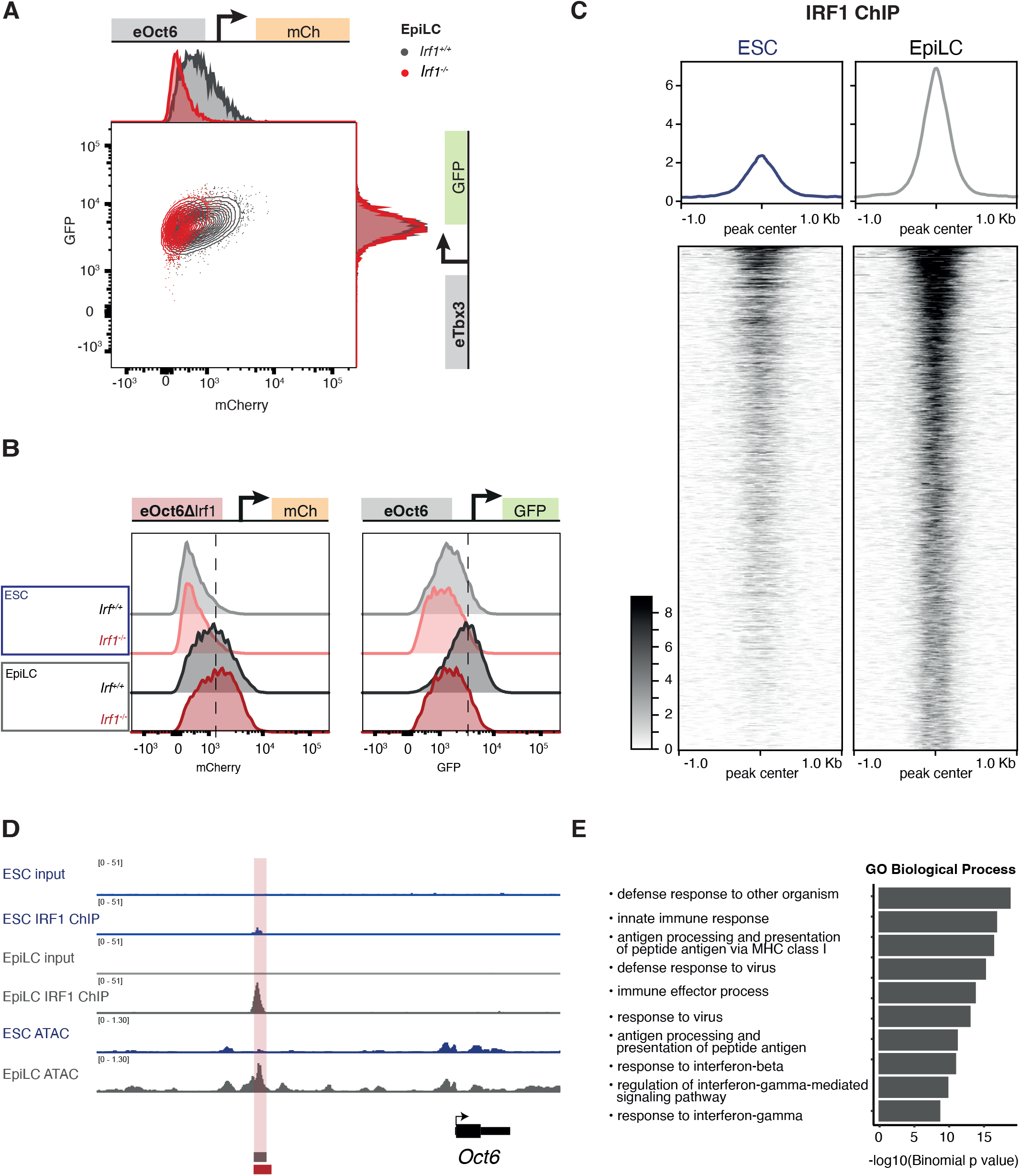
IRF1 regulates the *Oct6* enhancer and binds to genes involved in the innate immune response. **A** Representative flow cytometry profiles of reporter cell lines in EpiLC condition, *Irf1^+/+^* and *Irf1*^-/-^. See also Figure S3.1E. **B** Representative flow cytometry profiles of ESC and EpiLC cells, *Irf1^+/+^* and *Irf1*^-/-^ with *eOct6ΔIrf* and *eOct6* controlling mCherry and GFP, respectively. See also Figure S3.2A. **C** IRF1-ChIP-seq in ESC and EpiLC. Profiles and heatmaps of the consensus of all binding sites are shown. **D** Chromatin context of *Oct6* locus, with a called IRF1 peak highlighted. IRF1 ChIP-seq tracks and ATAC-seq tracks for ESC and EpiLCs are shown. The *eOct6* region used as reporter is marked with the lower red box. **E** GREAT-analysis of GO-term enrichment for IRF1 binding sites in ESC/EpiLC. Binomial p-values of top 10 GO biological processes are shown.

We then tested if e*Oct6* enhancer activity can be rescued by ectopic *Irf1* expression in the *Irf1* KO background. We decided to use the *eOct6* enhancer to drive *Irf1* expression since this allowed for differentiation induced IRF1 expression during the exit from naive pluripotency. IRF1 was already detected in ESCs at the protein level under rescue conditions **(Figure S3.1F**), most likely due to low enhancer activity in ESCs. This weak expression corresponded to an already increased e*Oct6* enhancer controlled mCherry signal in ESCs (**Figure S3.1G**). In EpiLCs, IRF1 expression levels were fully restored or even increased in comparison to endogenous levels (**Figure S3.1F**). This correlated to increased *eOct6* enhancer activity driving mCherry. Throughout all rescue conditions, *eTbx3* controlled GFP levels were not influenced and changed according to differentiation status (**Figure S3.1G**).

Since the *eTbx3* enhancer was not regulated by IRF1, we performed alkaline phosphatase staining to test whether *Irf1* KO cells displayed a general exit from naive pluripotency defect (**Figure S3.1H**). We differentiated cells for 72h to irreversibly exit naive pluripotency and plated them back under naive 2i+LIF conditions. Genotypes with exit from naive pluripotency defect show an increased fraction of cells still able to proliferate under 2i+LIF conditions. *Irf1* KO cells did not show increased colony formation and therefore are not impaired in the exit from naive pluripotency, consistent with the unimpaired regulation of the eTbx3 reporter.

In summary, we identified *Irf1* in our pooled CRISPR-KO screen as a regulator of the *eOct6* enhancer, which is a member of the formative pluripotency network. At the same time, loss of IRF1 does not affect the exit from naive pluripotency.

### IRF1 is directly activating the *Oct6* enhancer

The *eOct6* enhancer is less active in the *Irf1* KO cell lines, which could be due to direct regulation by IRF1 at this enhancer. Indeed, the *eOct6* enhancer itself contains a canonical IRF1 binding motif (gAAAgtGAAA). To test whether this motif is necessary for enhancer activity, we mutated the IRF1 motif (from here referred to as *eOct6-ΔIrf1*). We generated cell lines in WT and *Irf1* KO background with *eOct6-ΔIrf1* controlling mCherry expression. In addition, we included the wild type *eOct6* enhancer controlling GFP expression as an internal control **(Figure 3B, S3.2A**). As expected, the unmodified *eOct6* enhancer was activated in WT cells and to a lesser extent in *Irf1* KO cells. However, with the IRF1 binding motif mutated, the *eOct6-ΔIrf1* enhancer does not depend on the presence or absence of IRF1, as *Irf1* KO shows similar activation levels compared to WT cells. The IRF1 binding motif mutated *eOct6-ΔIrf1* enhancer is still increasing activity in differentiation into EpiLCs, stressing that IRF1 is not the sole regulator of *eOct6* activity, but that several factors are involved in the activation of the enhancer. Next we measured *Oct6* mRNA expression in RNA-seq of WT and *Irf1* KO ESC and EpiLC. *Oct6* mRNA upregulation in *Irf1* KO EpiLC is reduced compared to WT EpiLC, but not completely abolished (**Figure S3.2B**). This further suggests additional levels of regulation and redundancy. In conclusion, IRF1 directly regulates *eOct6* enhancer activity dependent on the IRF1 binding motif, but IRF1 is not the sole regulator of this enhancer.

### IRF1 binds to the *Oct6* enhancer and to genes involved in the innate immune response

Next, we asked at which chromatin sites endogenous IRF1 binds to in ESCs and EpiLCs. First, we validated IRF1 ChIP efficiency by qPCR. IRF1 ChIP recovers the promoter of *Gbp2* (**Figure S3.2C**), a known IRF1 bound site in macrophages (Ramsauer et al. 2007; Langlais, Barreiro, and Gros 2016). In addition, we also assayed two primer pairs at the *eOct6* enhancer locus, both showed enrichment in IRF1 Chip qPCR, confirming the direct binding of IRF1 to the *eOct6* enhancer locus (**Figure S3.2C**).

To test genome wide binding of IRF1, we sequenced libraries derived from IRF1 ChIP in WT and IRF1 KO for ESCs and EpiLC in replicates. Lack of peak calling by MACS2 for IRF1 KO samples confirmed specificity of the used antibody (**Figure S3.2D**). GO term enrichment across all stem cell IRF1 binding sites confirmed integration into biological processes such as defense response to other organisms and innate immunity processes (**Figure 3E**). Qualitatively, binding sites of IRF1 did not drastically change in differentiation (71 sites with FDR < 0.05). Quantitatively, IRF1 bound sites are weakly marked already in ESCs, and binding signal increases during differentiation into EpiLCs (**Figure 3C**). This is exemplified by the *Oct6* locus (**Figure 3D**). We compared ESC and EpiLC IRF1 binding sites to those in bone marrow derived macrophages (BMDMs), before and after interferon-ɣ stimulation (INFɣ) (Langlais, Barreiro, and Gros 2016) (**Figure S3.2E**). 64% of ESC/EpiLC IRF1 binding sites are also bound by IRF1 in BMDM, either stimulated or unstimulated with INFɣ (**Figure S3.2E**). We compared genomic annotations between the sites shared between BMDM and stem cells. Shared sites are located more often at promoter regions, while stem cell specific sites are located in intergenic regions, including enhancer sites (**Figure S3.2F**). A majority of IRF1 binding occurs at or close to promoters and seems to be conserved between stem cells and INFɣ stimulated BMDMs.

### IRF1 in pluripotency is a regulator of interferon stimulated genes

Next, we asked whether IRF1 has an effect on gene expression in differentiation from ESCs to EpiLCs. We performed RNA-seq using the QuantSeq protocol for WT and two independent *Irf1* KO cell lines under ESC and EpiLC conditions. Principal component analysis captured differentiation along PC1 and genotype specific effects along PC2 (**Figure 4A**). These results indicate that *Irf1* KO cells do not show drastically changed cell states in differentiation. To corroborate these findings, we focused on well-established pluripotency markers, expressed either in ESCs (such as *Klf4*, *Esrrb* and *Nanog*), or in EpiLCs (for example *Fgf5* and *Otx2*) and found little change in expression with the exception of *Oct6* as shown above (**Figure 4B**) (all shown factors p-value control vs *Irf1* KO > 0.05, besides Tbx3 (p-value = 0.031)). Previously, a list of 496 genes which are either positively or negatively associated with the core pluripotency markers was described (Naive Associated Genes, NAGs (Lackner et al. 2021)). NAGs are also correlated with pre- to post implantation expression changes in mice and macaque *in vivo*. As expected, NAGs show strong ESC and EpiLC specific expression, but no drastic change upon IRF1 deletion (**Figure S4A**). Therefore, IRF1 does not regulate the naïve or the formative gene regulatory networks.

**Figure 4.**
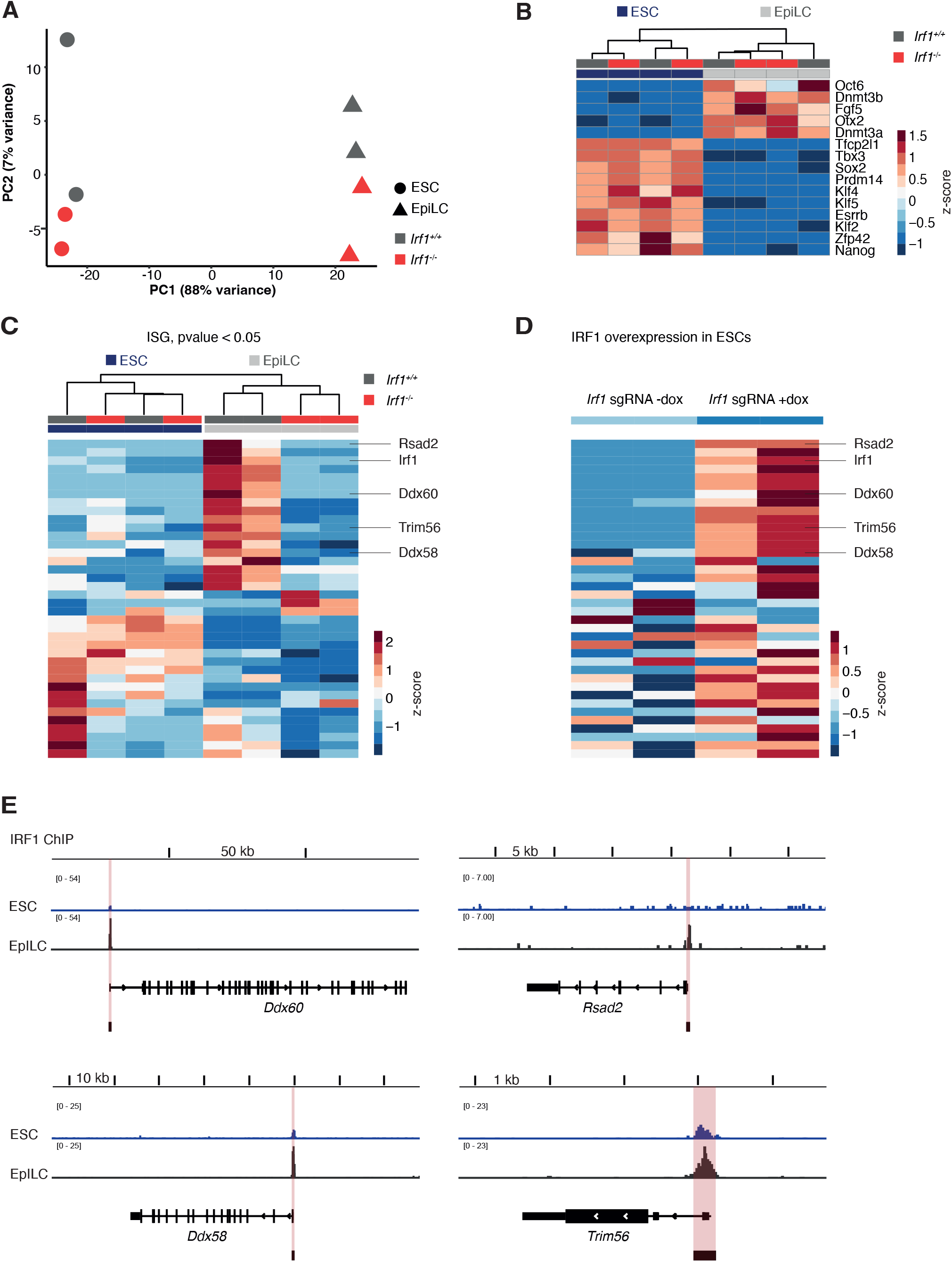
IRF1 regulates ISG expression in EpiLCs. A Principal component plot of *Irf1^+/+^* and *Irf1^-/-^* ESC and EpiLC, based on QuantSeq RNA data. B Expression changes of selected pluripotency markers in *Irf1^+/+^* and *Irf1^-/-^* ESC and EpiLC Quantseq RNA data. Data are shown as z-score. C Expression changes of ISG in *Irf1^+/+^* and *Irf1^-/-^* ESC and EpiLC Quantseq RNA data. Shown are ISG differentially expressed (p-value< 0.05) between *Irf1^+/+^* and *Irf1^-/-^* EpiLCs. Data are shown as z-score. See also Figure S4C. D Expression changes of the same genes as in Figure 4C in dox-inducible SunTag-based IRF1 overexpression. Data are shown as z-score. Selected gene names are indicated. E Chromatin loci with IRF1 binding profiles for genes selected in Figure 4D. Called IRF1 peaks are highlighted by red boxes.

*Irf1* is a member of a group of genes previously classified as ISGs. Strikingly, a selection of ISGs are intrinsically expressed in stem cells of several species, even in the absence of viral infection (Wu et al. 2018). At the same time, stem cells are resistant against viral infection. In differentiation to somatic cell types, this intrinsic ISG expression and with it the viral protection is lost (Wu et al. 2018). We first confirmed high expression levels of ISGs (as defined in Wu et al. 2018) in our ESC and EpiLC RNA seq data (**Figure S4B**). As expected, a subset of ISGs was expressed in ESCs and in EpiLCs, however, we also observed changes in the cell state specific expression of ISGs (**Figure S4B**).

Given the role of IRF1 as a core component of the interferon-ɣ response, we asked if ISG expression is influenced by IRF1 in pluripotent cells. We tested which ISGs are differentially expressed in *Irf1* KO vs WT in EpiLCs (**Figure S4C**). Out of this subset of ISGs, *Irf1* KO most strikingly affects genes which under normal differentiation conditions are upregulated (**Figure 4C**).

We asked whether these genes are under direct control of IRF1 and established an IRF1 overexpression system in ESCs using an inducible CRISPR-ON system (Heurtier et al. 2019). CRISPR guide RNA are constitutively expressed, upon doxycycline (dox) addition, dCAS9 coupled to 10x GCN4 is induced, which in turn recruits several molecules of scFv-linked VP64, a potent activator. We used an sgRNA targeting the *Irf1* promoter and validated expression of full length IRF1 protein 48 hours after dox induction in ESC (**Figure S4D**). We performed QuantSeq to detect RNA expression changes after IRF1 overexpression. As a control, we included dox induction of dCAS9 in absence of sgRNA expression. Activity of untargeted dCAS9 had a strong effect on gene expression, potentially because of off-target effects (**Figure S4E**)(Hsu et al. 2013). As expected, IRF1 expression was upregulated when sgRNA directed to the promoter of *Irf1* was used under dox induction conditions (**Figure S4F**). Expression of NAGs was not influenced by IRF1 overexpression, but rather by non-targeted dCAS9 presence (**Figure S4G**, left panel). In contrast, ISG and specifically those ISG that were also misregulated in *Irf1* KO, showed upregulation in IRF1 overexpression (**Figure S4G**, right panel**, Figure 4D**).

Most of these genes were annotated as genes closest to IRF1 ChIPseq peaks and were also bound by IRF1 in BMDMs (**Figure 4E**). Regulated genes include core components of the interferon response: For example, *Ddx58/Rig-I* and its ligand-specific sentinel *Ddx60* (Oshiumi et al. 2015) showed increased IRF1 binding at the promoter in differentiation and subsequent upregulation during the transition from ESC to EpiLCs. During overexpression of IRF in ESCS, these factors are also upregulated, establishing both genes as direct targets of IRF1 in the exit from naive pluripotency. DDX58/RIG-I is known as a sensor of viral RNA (Kell and Gale 2015). Other examples include *Trim56* and *Rsad2/Viperin*, which are also bound and regulated by IRF1 and are well-established components of innate immune response. Taken together, ISGs directly regulated by IRF1 are conserved between early embryonic and somatic cell types.

### *Irf1* is activated by the formative pluripotency gene regulatory network

ISGs are expressed in stem cells in the absence of exogenous stimuli, but the mechanism establishing this intrinsic expression is unknown. We analyzed the chromatin environment surrounding the *Irf1* gene to identify potential regulatory regions that might explain the expression changes. 10 KB upstream of *Irf1* we identified a putative enhancer region. In ESCs, this element is not strongly marked by active enhancer marks. However, in EpiLCs this element is marked by OCT4, P300, the formative transcription factor OTX2 and the active histone modifications H3K27ac and H3K4me1 (**Figure 5A**). Importantly, this putative enhancer is specific for the ESC to EpiLC transition since it was not marked by H3K27ac in unstimulated and IFN-ɣ exposed BMDMs. In contrast, in BMDMs a region 6 kb upstream of *Irf1* is marked by increasing H3K27ac after IFN stimulation. This suggests a cell type specific regulation mechanism for *Irf1* in stem cells.

**Figure 5.**
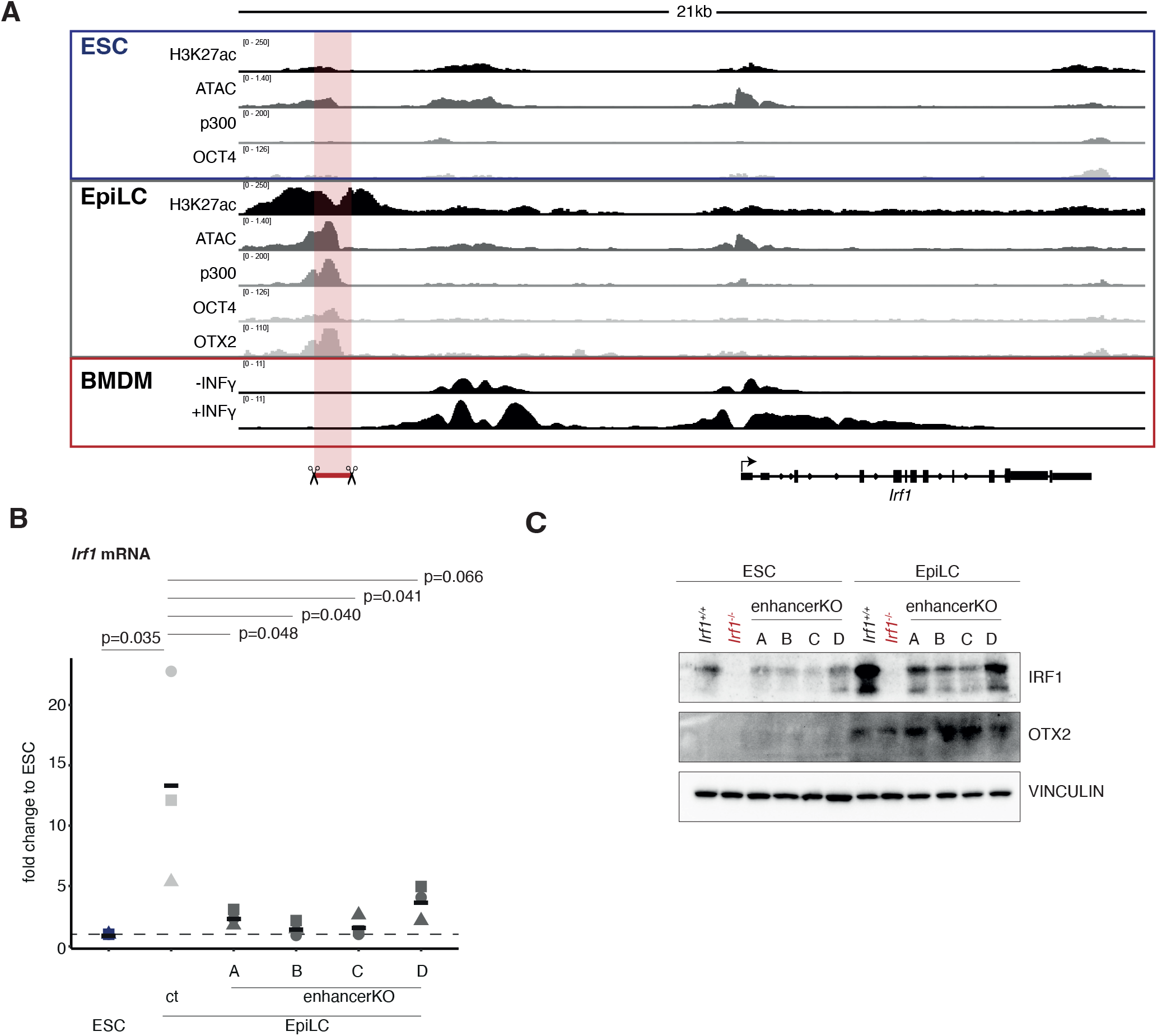
*Irf1* expression is controlled by the formative gene regulatory network. **A** Chromatin context of *Irf1* in ESCs, EpiLC and BMDMs with and without INFɣ. ESC/EpiLC ChIP data was generated by (Buecker, 2014), BMDM data by (Langlais, Barreiro, and Gros 2016). An putative enhancer which was knocked out is indicated. **B** RT-qPCR analysis of *Irf1* mRNA in *Irf1^+/+^* and *Irf1* enhancer KO in differentiation. Fold change is normalized against *Rpl13a* housekeeping mRNA expression and is calculated against ESC for each indicated cell line. This is shown as baseline with the dashed line at fold change = 1. Statistical tests were performed as homoscedastic one sided t-tests. **C** Western Blot analysis of *Irf1^+/+^*, *Irf1^-/-^* and *Irf1* enhancer KO ESC and EpiLC, probed with antibodies against IRF1, formative marker OTX2 and VINCULIN as loading control.

To analyze whether this putative enhancer element indeed activates the expression of IRF1, we deleted this enhancer region. We generated four independent enhancer KO cell lines and validated the absence of the enhancer region by genotyping PCRs and Sanger sequencing of the PCR products **(Figure S5A, S5B**). We then tested if endogenous IRF1 is upregulated to the same extent as in WT cells during differentiation into EpiLCs. Enhancer KO lines showed significantly reduced *Irf1* mRNA levels (**Figure 5B**). We also confirmed reductions of IRF1 protein levels by Western Blot (**Figure 5C**). As suggested from RNA sequencing data, the formative marker OTX2 is not influenced by the absence or reduction of IRF1 in EpiLCs. This enhancer region of *Irf1* connects the pluripotency network to the expression of IRF1.

In summary, IRF1 connects the formative pluripotency network and the regulation of ISG in pluripotent stem cells.

## Discussion

In this study, we performed a pooled CRISPR-KO screen to identify factors whose deletion impair the activation of the formative pluripotency gene regulatory network. For this, we used expression dynamics of a dual reporter system to identify cells which are showing expected behavior - are unimpaired in differentiation - or those which are impaired in differentiation. In contrast to CRISPR-KO screens which for example depend on proliferation of cells over prolonged time frames, our screening set-up is more sensitive to fluctuation due to noise. We counteracted these issues by focusing on a smaller library selected for mostly nuclear factors, which could miss important factors like metabolic regulators (Moussaieff et al. 2015). Nevertheless, we identified several factors such as *Fgfr1* and *Zic3* which have already been studied in the context of the exit from naive pluripotency, stressing the validity of this approach (Molotkov et al. 2017; Yang et al. 2019).

We infected cells with a low multiplicity of infection (MOI) to ensure single KO of one gene per cell. However, recent studies have shown the markable redundancy ensuring differentiation from naive to formative pluripotency: KO of single factors delays, but not completely abolishes differentiation (Lackner et al. 2021). Differentiation is only abolished by simultaneous genetic deletion of three complementary drivers (Kalkan et al. 2019). Furthermore, the naive pluripotency network is supported by functionally overlapping factors of the same orphan nuclear receptor family (Festuccia et al. 2021). We therefore speculate that cooperativity and redundancy between two or more factors are ensuring proper execution of this cell state transition, and that screening approaches should be adapted to this in the future.

We identified *Irf1* in our CRISPR-KO screen as influencing the reporter activity of an enhancer region of *Oct6*, which we used as proxy for the establishment of formative pluripotency. However, IRF1 deletion does not drastically change gene expression of other formative markers such as *Otx2*, *Fgf5* and *Dnmt3a/b*, indicating that observed change in reporter activity is not strictly a readout of cell state. IRF1 is directly interacting with the chosen enhancer region through a canonical IRF1 chromatin binding motif. *Oct6* gene expression is reduced in the absence of IRF1. *Oct6* belongs to the family of POU TFs that share a highly similar DNA recognition motif. It was shown that OCT6 can replace OCT4 during reprogramming into human iPS cells due to the similarity in DNA motif binding (Kim et al. 2020). OCT6 and OCT4 occupy many of the same sites in Epiblast stem cells (EpiSCs) and it is therefore not surprising that lower levels of OCT6 have only minor effects in the exit from naive pluripotency (Matsuda et al. 2017). Furthermore, deletion of *Oct6* in the mouse does not severely impacts embryogenesis, again probably due to the functional redundancy among different members of the POU transcription factor family (Kim et al. 2021), (Bermingham et al. 1996).

Redundancy seems to be present not just in naive pluripotency, but also among many formative pluripotency regulators: the transcription factor OTX2 is sufficient to drive gene expression changes associated with formative pluripotency, but its deletion has only limited effect on the expression of formative genes (Buecker et al. 2014; Yang et al. 2014). Misregulation of OCT6 could therefore also be compensated by other members of the formative gene regulatory network. While OCT6 has been studied mostly in the context of development and neurogenesis, it might have additional roles upon interferon stimulation. OCT6 itself is upregulated by interferons in macrophages and other cell types, however, its role in interferon signaling is unclear (Hofmann et al. 2010).

Importantly, *Irf1* itself is directly regulated by the formative pluripotency network through an enhancer which is marked by activating chromatin marks and OCT4 and OTX2 in formative EpiLC. The same enhancer does not get activated in stimulated BMDMs and therefore seems to be stem cell specific (Lara-Astiaso et al. 2014). This direct regulation could explain the expression of specific ISGs in stem cells. ISG expression in early embryonic tissues, iPS or also somatic stem cells is conserved in mouse, pig, chimpanzee and human (Wu et al. 2018; Shi et al. 2020). This expression is independent from viral stimulation and was previously only described as intrinsic (Wu et al. 2018) or as a consequence of endogenous retroviral expression observed in hypomethylated stem cells (Grow et al. 2015). Interestingly, Wu and colleagues showed that ISG expression in stem cells can protect these cells from viral infection. We show here that the formative pluripotency network itself is controlling the expression of a subset of ISGs through upregulation of IRF1 and might thereby protect this stage from viral infection. In turn, this connection through IRF1 signals back to the pluripotency network through upregulation of OCT6, albeit other factors might be in place to ensure robust pluripotency. Robustness against viral infection could be an important property of proliferating stem cells and is therefore controlled by the specific gene regulatory network that safeguards the stem cell state.

## Acknowledgement

We thank all members of the Buecker lab for discussions and feedback throughout the project, Martin Leeb, Thomas Decker and Gijs Versteeg for critical feedback and discussion on the manuscript and the BioOptics-FACS and BioOptics-Light Microscopy facility at Max Perutz Labs.

The ATAC-seq, ChIP-seq and QuantSeq was performed by the Next Generation Sequencing Facility at Vienna BioCenter Core Facilities (VBCF), member of the Vienna BioCenter (VBC), Austria.

This work was supported by the Austrian Science Fund FWF (P30599 and P34123 to C.B. and W1261 DK SMICH) and a Uni:Docs fellowship from the University of Vienna to M.R.

## Author contributions

C. B and M.R. conceived and designed the study. M.R., S.S. and M.H. generated and validated cell lines and carried out experiments. S.F. generated and validated enhancer KO cell lines. M.R. carried out the screen. M.R. and G.V. analyzed screening data, with input from U.E.. M.R. analyzed all sequencing data. M.R. and C.B. wrote the manuscript, with input from all co-authors. C.B. supervised the project.

## Methods

### ESC maintenance and differentiation conditions

Murine embryonic stem cells were cultured and differentiated as described in (Thomas et al. 2021). For maintenance, cells were grown on CELLSTARÒ 6/12-wells coated with first poly-L-ornithine hydrobromide (6 µg/ml in PBS, 1h at 37 °C, sigma P4638) and then laminin (1.2 µg/ml in PBS, 1 h at 37°C, sigma L2020). For differentiation and viral infection, plates were coated with fibronectin (Human Plasma Fibronectin Purified Protein, Sigma-Aldrich, 5µg/ml in PBS, 1h at RT).

Cells were cultured in base medium HyClone DMEM/F12 without Hepes (Cytiva) with 4 mg/mL AlbuMAX^™^ II Lipid-Rich Bovine Serum Albumin (GIBCO^™^), 1x MACS NeuroBrew-21 with Vitamin A (Miltenyi Biotec), 1x MEM NEAA (GIBCO^™^), 50 U/mL Penicillin-Streptomycin (GIBCO^™^), 1 mM Sodium Pyruvate (GIBCO^™^) and 1x 2-Mercaptoethanol (GIBCO^™^). For 2i+LIF culturing conditions, base medium was supplemented with with 3.3 mM CHIR-99021 (Selleckchem), 0.8 mM PD0325901 (Selleckchem) and 10 ng/mL hLIF (provided by the VBCF Protein Technologies Facility, https://www.viennabiocenter.org/facilities/). For differentiation, base medium was supplemented with 12 mg/mL Recombinant Human FGF-basic (PEPROTECH) and KnockOut^™^ Serum Replacement (1:100, GIBCOTM).

For splitting, cells were treated with 1x Trypsin-EDTA solution (sigma T3924) at 37 °C until cells detached. Trypsinization was stopped with 2i+LIF medium with 10% FSC (sigma F7524), cells were harvested by 300g for 3 min, resuspended in 2i+LIF and seeded in appropriate ratios.

For differentiation, cells were seeded a day prior on fibronectin coated plates in 2i+LIF medium (100k per 12 well, 200k per 6 well). The next day, differentiation was started by removing the medium, 2 washes of the attached cells in PBS, and addition of differentiation medium. Cells were collected as EpiLCs after 48 hours.

### Generation of reporter and screening cell lines

Cells used for the dual fluorescent reporter system and CRISPR screening were based on an Otx2^flox/-^; R26^CreER/+^ ESC cell line (Acampora, 2013). Otx2 heterozygous expression does not impair ESC to EpiLC differentiation (Acampora, 2013). pB-transposase, pB-eOct6-mCherry-Puro and pB-eTbx3-dGFP-Hygro were lipofectamine transfected. After selection, two clonal cell lines with clear distinction of ESC/EpiLC states in FACS analysis were selected. Lentivirus containing Cas9-Blasticidin was used for constitutive Cas9 expression (kind gift from Ulrich Elling). Neomycin resistance derived from the original present LacZ allele was removed by CRISPR KO.

For pB-eOct6ΔIrf1-mCherry-Puro, the mutated IRF1 binding sites were placed on overlapping primers, the Oct6 enhancer was PCR amplified and the plasmid constructed with three fragment Gibson Assembly. pB-transposase, pB-eOct6ΔIrf1-mCherry-Puro and pB-eOct6-mCherry-Neo were co-transfected into ESCs with lipofectamine.

### Generation of KO and rescue cell lines

Knockouts of coding sequences or enhancer regions were performed with dual CRISPR guide KO. For this, CRISPR guides were inserted into pX330-U6-Chimeric_noCas9 (for cell lines already expressing Cas9) or pX330-U6-Chimeric_BB_CBh_hSpCas9 using BbsI (NEB) directed cloning. Guide plasmid combinations and substoichiometric amounts of dsRed plasmid were cotransfected by Lipofectamine® 2000 (Invitrogen) transfection (375 ng each guide plasmid, 50 ng dsRed, 5 µl lipofectamine in total 100 µl DMEM/F12) into cells seeded on 12-well plates one day prior. 2-3 days after transfection, dsRed+ cells as proxy for guide transfection were FACS-sorted (BD FACSMelody Cell Sorter) as single cells on 96 well plates coated with fibronectin. Successful genome editing was confirmed by genotyping PCRs and Western Blotting where applicable.

For rescue of Irf1 expression we cloned pB expression constructs containing *Irf1* cDNA under control of the Oct6 enhancer. This resulted in low IRF1 expression levels in ESCs and upregulation of IRF1 upon differentiation. Transfection of pB-Oct6enh-Irf1-Neo was performed with lipofectamine transfection as described for reporter constructs, albeit with lower DNA amounts (250 ng pB plasmid). After neomycin selection (400 µg/ml, G415 Disulfate solution, sigma G8168-10ML), pools of transfected cells were differentiated and analyzed by FACS and Western Blotting.

### IRF1 Overexpression with SunTag system

Overexpression of Irf1 from the endogenous locus was based on a SunTag overexpression system published in Heurtier et al, 2019. Following plasmids were ordered from Addgene: 121121 PB-gRNA-Puro, 121119 PB-TetON-dual-SunTag-Hygro, 20910 PB-CA-rtTA Adv. 10 µg PB Transposase, 500 ng PB-TetON-dual-SunTag-Hygro and 500 ng PB-CA-rtTA were transfected into R1 ESCs by electroporation (500 µF, 240 V, 4mm). Cells were treated with doxycycline (1µg/ml) and hygromycin (400 µg/ml). After 5 days, BFP+/GFP+ double positive cells were selected by FACS. Cells were grown in the absence of doxycycline and hygromycin and BFP-/GFP- double negative cells were selected by FACS. Single clones were selected and tested by FACS and Western Blotting against CAS9 to select for response to doxycycline treatment.

Guides targeting the Irf1 promoter were designed in benchling, which is based on (Hsu et al. 2013) and cloned into PB-gRNA-Puro by BbsI directed cloning. The guide plasmid and PB transposase were lipofectamin transfected into parental SunTag cell lines and selected by puromycin (2 µg/ml, Invivogen) treatment. *Irf1* upregulation after dox treatment was confirmed by qPCR and Western Blotting.

### CRISPR screening

CRISPR screening was based on Michlits et al. 2017, but without using clonal dilution steps. DNA pools of indicated subpools were combined according to guide number and CaCl/Hepes (Thermo Fischer Scientific 7365-45-9) transfected into PlatE cells, including helper plasmid. 24 hours after transfection, medium was exchanged to 2i+LIF. Virus containing 2i+LIF was harvested in two batches over the next 36 hours. Both batches were pooled and frozen. MOI was determined by determining infection efficiency without selection.

### Execution of CRISPR screen

The screen was performed with two independent replicates with different screening cell lines. To reach sufficient coverage of the library, we infected 100 mio cells at a MOI of 0.3 by seeding 7.5 mio cells per 15 cm dish coated with fibronectin. Starting library representation was accessed by infecting cells without Cas9 (R1) in parallel. In addition, no selection, kill and no guide library control plates were prepared. Usage of fibronectin coated plates allows to omit polybrene addition during infection. As soon as cells were attached (2 h), virus was added to the cells without additives or medium exchanges (day1). The next day, selection with neomycin was started and continued for 2 days. From now on, each day cells were either splitted or medium was exchanged. On day 4, the kill control was dead and 30 mio library infected cells were seeded on 15 cm PLOL plates with 7.5 mio per plate without selection. Empty library controls were carried throughout the screen. On day 6 and 8, again 30 mio cells were seeded without selection, allowing time for depletion of essential genes. On day 10, 60 mio cells were harvested and analyzed later as samples representing genes essential for ESC maintenance. For differentiation, 30 mio cells were seeded on fibronectin coated dishes (7.5 mio per 15 cm dish) Additionally, extra dishes as 2i+LIF control and no guide library controls were seeded. On day 11, differentiation was induced by 2+iLIF medium removal, 2 washed in PBS and addition of differentiation medium. Starting timepoints for different plates were staggered to allow FACS analysis after 48 of differentiation for each plate.

On day 13, cells were harvested and sorted by BD FACSAria (BD Biosciences). For this, gates were set up using no guide library control cells, which were processed in parallel. At least 60 mio cells were sorted in the unimpaired gate and all possible cells in the impaired gate (N1 = 64.7 to 10.3 mio, N2= 71.5 to 5.6 mio cells). In addition, 60 mio cells for essential ESC and 60 mio for unsorted EpiLCs were harvested for each condition. The screen was performed as replicate using two independently generated cell lines.

### gDNA isolation, PCR amplification and NGS

CRISPR KO libraries were generated from collected cells as described by (Michlits et al. 2017), scaled according to the cell numbers used.

In brief, cell pellets were resuspended in SDS-lysis buffer and incubated at 55 °C overnight to lyse cells, followed by RNaseA treatment. Protein was precipitated by NaCl precipitation, the supernatant was then purified with phenol/chloroform. DNA was precipitated with isopropanol and dissolved in TE buffer. DNA was PacI digested for a total of 48 h and size selected with SpeedBeads™ (Millipore Sigma) in two rounds to enrich fragments < 2 kb.

CRISPR-UMI cassettes were PCR amplified and multiplexed with 25 cycles of 1:1 KlenTag/Phusion PCR (Jena Bioscience and in house generated) using primers specified in (Michlits et al. 2017). Reactions were purified using the NucleoSpin PCR and Gel Purification Kit (Macherey-Nagel). Samples were pooled after qPCR quantification, purified from 1.5% agarose and quantified with NEBNext Library Quant Kit for Illumina (NEB). Sequencing was performed on Illumina HiSeqV4 SR50.

### RT-qPCR analysis

ESC and EpiLC were directly lysed on the plate using pepGOLD TriFast^™^ reagent (Peqlab) according to manufacturer’s instructions. 500 ng of RNA were reversed transcribed to cDNA with the SensiFast^™^ cDNA Synthesis kit (Bioline) according to manufacturer’s instructions. For qPCR, cDNA was 1:5 diluted and 0.5 µl were used per 10µl qPCR reaction with SensiFAST^™^ SYBR^(R)^ No-ROX kit (Bioline) and 125 nM forward and reverse primer. Irf1 specific primers were based on (Platanitis et al. 2019).

qPCRs were analysed by ΔΔCT and normalized to ESCs samples of the corresponding cell lines. Statistical tests were performed as homoscedastic one sided t-tests.

### FACS analysis

ESC and EpiLC were harvested by trypsin treatment, which was stopped by 1:1 addition of 10% FSC. Samples were strained through 5 ml Polystyrene Round-Bottom Tubes with Cell-Strainer Caps (Falcon). For analytical purposes, the BD Fortessa was used, for cell sorting in high-throughput BD FACSAria or in low-throughput BD FACSMelody. Single cells were gated according to FFW and SSW scattering.

### QuantSeq

RNA QuantSeq was prepared according to manufacturer’s instructions (Lexogen 3’ mRNA Seq Library Prep Kit). 500 ng RNA was used as starting material. qPCRs were used to multiplex samples and final libraries were quantified by NEBNext Library Quant Kit for Ilumina (#E7630S).

### Chromatin Immunoprecipitation

ChIP with Irf1 was performed as described in (Thomas et al. 2021). In brief, 3 mio cells were seeded on 15 cm fibronectin coated dishes, medium exchanged to 2i+LIF or differentiation the next day and cells harvested after 48 hours.

For harvesting, cells were crosslinked in 1 % formaldehyde in PBS for 10 min. Crosslinking was quenched in 0.125 M glycine. From now on samples were kept at 4 °C. Fixed cells were washed on the plate twice with PBS and scraped off in 0.01 % Triton in PBS. Cells were harvested by 500g 5 min centrifugation and flash frozen in liquid nitrogen. Cell pellets were resuspended in 5 ml LB1 (50 mM Hepes pH 7.5, 140 mM NaCl, 1 mM EDTA, 10% glycerol, 0.5% NP-40, 0.25% TX-100, 1 mM PMSF, 1x cOmpleteTM Protease Inhibitor Cocktail (Roche) by rotating vertically for 10 min at 4°C and harvested (5 min, 1350g, 4°C). The pellet was resuspended in 5 ml LB2 (10 mM Tris pH 8. 200 mM NaCl, 1 mM EDTA, 0.5 mM EGTA1, mM PMSF,1x cOmpleteTM Protease Inhibitor Cocktail (Roche)), 10 min vertical rotation, room temperature. Centrifugation was repeated and pellets were suspended in 1.5 ml LB3 (10 mM Tris-HCl pH 8, 100 mM NaCl, 1 mM EDTA 0.5 mM EGTA, 0.1 % Na-deoxycholate, 0.5% N-lauroylsarcosine, 1 mM PMSF, 1x cOmpleteTM Protease Inhibitor Cocktail (Roche)) and 200 µl sonification beads (diagenode) in Bioruptor(R) Pico Tubes (diagenode). Sonification was performed for 13 cycles 30s on /45s off. Supernatant, but not the beads were transferred to fresh tubes and cellular debris was pelleted at 16000g at 4 °C. 1.1 ml were transferred to fresh tubes and 110 µl 10% triton to final 1 % were added. 50µl was saved as input and 1 ml was used for ChIP.

Antibody precipitation was performed with 5 µl IRF1 antibody overnight at 4°C with vertical rotation. Next day, Dynabeads protein G for Immunoprecipitation (Thermo Fisher Scientific) were used. 100 µl beads were washed in block solution (0.5% BSA in PBS) and incubated with chromatin/antibody solutions for at least 2 hours. Bound beads were washed 5 times in cold RIPA wash buffer (50 mM Hepes pH 7.5, 500 mM LiCl, 1 mM EDTA, 1 % NP-40, 0.7% Na-Deoxycholate), followed by 3 washes in TE + 50 mM NaCl. Bound fractions were eluted in 210 µl elution buffer (50 mM Tris pH 8.0, 10 mM EDTA, 1 % SDS) for 15 min at 65°C. Supernatant was removed from the beads. Input samples were diluted with 3 volumes of elution buffer. Input and ChIP samples were decrosslinked at 65 °C overnight.

One volume of TE was added to all samples as well as RNase A (0.2mg/ml final) and incubated at 37 °C for 2 hours. Salt concentration was adjusted to final 5.25 CaCl2 with 300 mM CaCl2 in 10 mM Tris pH 8.0 and Proteinase K added to final 0.2 mg/ml. Digestion was performed at 55 °C for 30 min. DNA was phenol-chloroform extracted in Phase Lock Gel^™^ tubes (Quantabio), ethanol precipitated and dissolved in H20.

ChIP was checked by performing qPCRs and calculating recovery of input (%).

Sequencing libraries were generated with sparQ DNA Library Prep Kit (Quanta Bio #95191-096) according to manufacturer’s instruction and using AMPure XP beads. Adapter contamination was cleaned up with an additional round of AMPure XP bead purification. Quality of the libraries were checked by Bioanalyser (Agilent) and final concentrations were determined with the NEB library Quant Kit.

### ATACseq

ATACseq was performed for ESC and EpiLCs. Nuclei were isolated as described by 10X Genomics, “Nuclei Isolation for Single Cell Multiome ATAC + Gene Expression Sequencing (Demonstrated Protocol CG000364)”. 50000 nuclei were incubated with Tn5 transposase (Illumina #15027865) in TD buffer at 37°C, 30, 600rpm. DNA was purified with Qiagen MinElute columns and in-house produced MBSpure beads (Ampure XP alternative). For this 13 µl eluted DNA were mixed with 25µl 8M Guanidinium hydrochlorid solution pH 8.5. After 5 min, binding buffer (20% PEG 8000, 2.5 M NaCl, 10 mM Tris-HCl pH8, 1mM EDTA, 0.05% Tween 20) and 30 µl MBSpure beads were added and incubated for 10 min. Beads were bound by magnet and washed twice in 150 µl 80% EtOH. Beads were dried for 30s and resuspended in 50 µl H20. 25 µl binding buffer was added, incubated for 5 min and magnetized. Supernatant was transferred to fresh wells. 50 µl AMPure beads were added, washed twice in EtOH, and final DNA was eluted in 20 µl H20. Libraries were amplified with i5/i7 Nextera primer index mix and Q5 PCR Master Mix (NEB #M0492L): 5 min 72°C, 1 min 98°C, 6x[10s 98°C, 30s 65°C, 30s 72°C]. Libraries were again purified by AMPure beads, quality controlled by Bioanalyser and quantified by NEBNext Library Quant Kit for Illumina (#E7630S).

### Alkaline Phosphatase Staining

For AP staining, cells were seeded in different densities on gelatine coated plates. The next day, differentiation was performed for 72 hours, controls were kept in 2i+LIF medium. All cells were placed back in 2i+LIF medium. After 2-4 day, colonies were stained with the Alkaline Phosphatase Detection Kit (Sigma-Aldrich) according to manufacturer’s instructions.

### Immunofluorescence

10k cells were seeded per chamber of a µ-Slide 8 Well Chambered Coverslip (ibidi). Cells were differentiated for 48 hours, washed in PBS and fixed with 4% PFA for 15 min at RT. Cells were washed 3x in PBST (0.1 % Tween in PBS). Permeabilization was performed in 0.1% Triton-X in PBS for 10 min at RT. Cells were washed 3x in PBST and blocked in 5% BSA in PBST for 30 min at 4 °C. Primary antibody was diluted in blocking buffer and incubated overnight at 4 °C. Cells were washed 5x in PBST and incubated with secondary antibody in blocking buffer for 1 hour at RT. Cells were washed 3x in PBST, followed by 2x PBS washes. Nuclei were stained with 20 ng/ml DAPI (Sigma, D9542) for 10 min. Cells were washed 3x in PBS and stored in PBS at 4 °C until imaging. Imaging was performed on a Zeiss Axio Oberserver Z1 microscope with 63X oil objective. Images were processed using ImageJ.

### Western Blot Analysis

For Western Blot analysis, cells were harvested by trypsin treatment. The cell pellet was washed once in PBS, after supernatant removal dry pellets were frozen. Cells were lysed in 1X RIPA lysis buffer (Merck, 20-188), including 1x cOmpleteTM Protease Inhibitor Cocktail (Roche) and phosphatase inhibitors (1 mM NaF, 20 mM ꞵ-glycerophosphate, 1 mM Na3VO4) for 1 hour on ice. Cellular debris was removed by centrifugation (16000g, 10 min, 4 °C). Protein was quantified using Bio-Rad Protein Assay (5000006). 25 µg protein per sample were separated on 10% Tris-gylcine SDS polyacrylamid electrophoresis. Wet-Blotting was performed onto PVDF Transfer Membranes (Therome Scientific #88518). Membranes were blocked in 5 % milk in PBST. Primary antibodies were incubated overnight at 4°C, secondary for 2 hours at RT. HRP coupled secondary antibodies were used for detecting with GE Healthcare LS ECL Select WB detection reagent. Images were acquired on a Chemidoc Touch (Bio-Rad).

### Data analysis

For CRISPR KO screening, reads were aligned to the library using custom scripts as published in (Michlits et al. 2017). Starting from count tables, analysis was performed with MAGeCK 0.5.9.2 and R 4.0.4. Plots were generated with ggplot2 (3.3.5)

QuantSeq RNA seq data was processed following Lexogen’s standard pipeline on bluebee. Adapter contamination was removed with bbmap, mapped using STAR and counted with HTScount. Downstream analysis was performed in R using DESeq2 (1.30.1) and pheatmap (1.0.12).

For ChIP and ATAC analysis, the respective Nextflow/21-04.1 nf-core pipelines were used with mm10 as reference genome. Profiles and heatmaps were generated with deeptools 3.5.1. ChIP data was analyzed with ChIPpeakAnno (3.24.2) and GO term where determined with GREAT (4.0.4) analysis (McLean et al. 2010).

ISGs were defined based on (Wu et al. 2018).

### plasmids

| plasmid | source |
| --- | --- |
| pB-eOct6-mCherry-Puro | this study |
| pB-e/Oct6--GFP--Neo | (Thomas et al. 2021) |
| pB-eOct6ΔIrf1motif-mCherry-Puro | this study |
| pB-eTbx3-dGDP-Hygro | this study |
| pX330-U6-Chimeric_BB_CBh_hSpCas9 | Addgene Plasmid #42230 |
| pX330-U6-Chimeric_noCas9 | this study |
| pB-transposase | System Biosciences Cat#PB210PA-1 |
| PB-gRNA-Puro | Addgene Plasmid #121121 |
| PB-TetON-dual-SunTag-Hygro | Addgene Plasmid #121119 |
| PB-CA-rtTA Adv | Addgene Plasmid #20910 |
| pB-Oct6-Irf1-Neo | this study |

### Primer

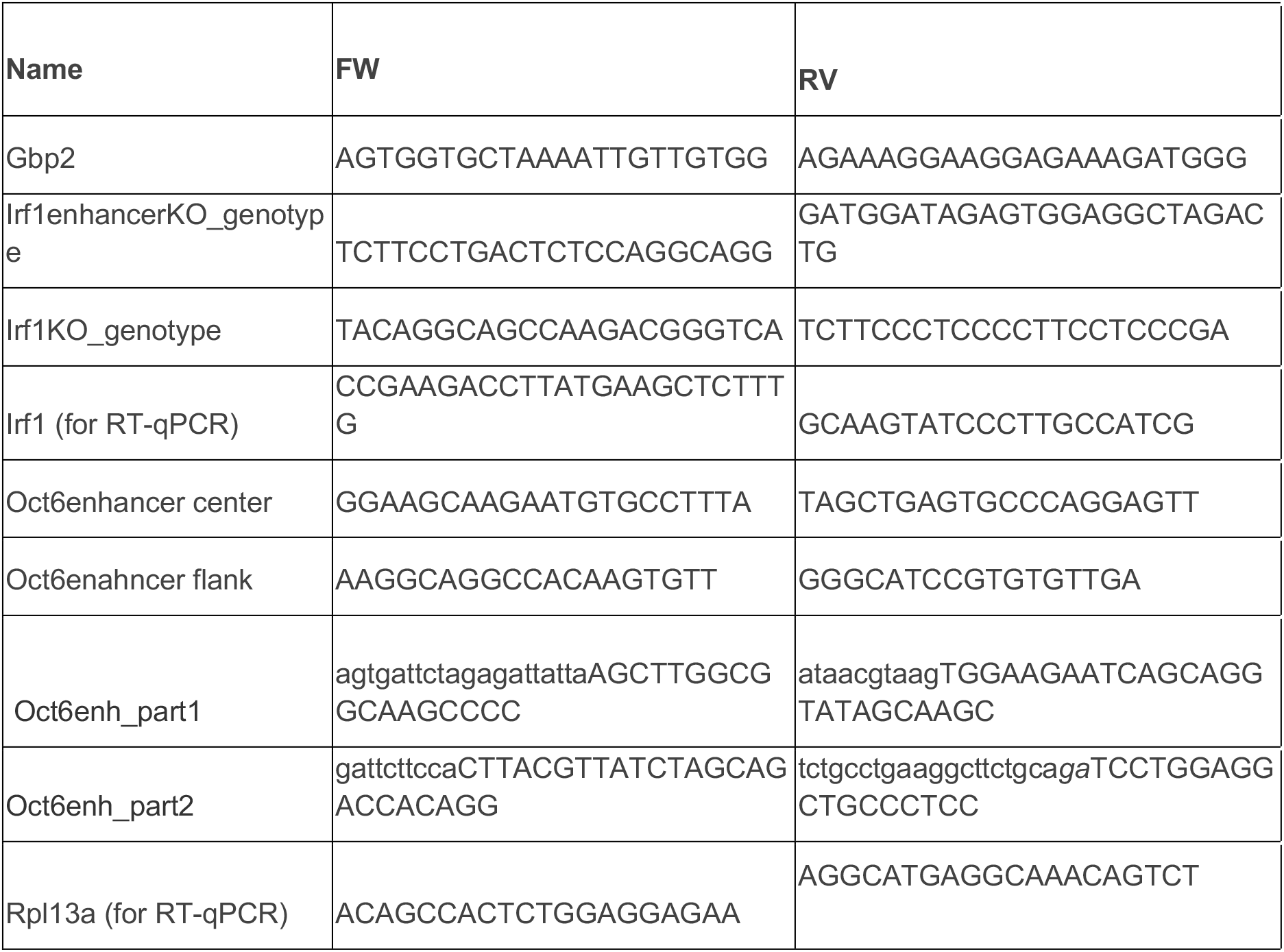

### guides

Sequences include sites for BbsI directed cloning, forward and reverse strands are annealed.

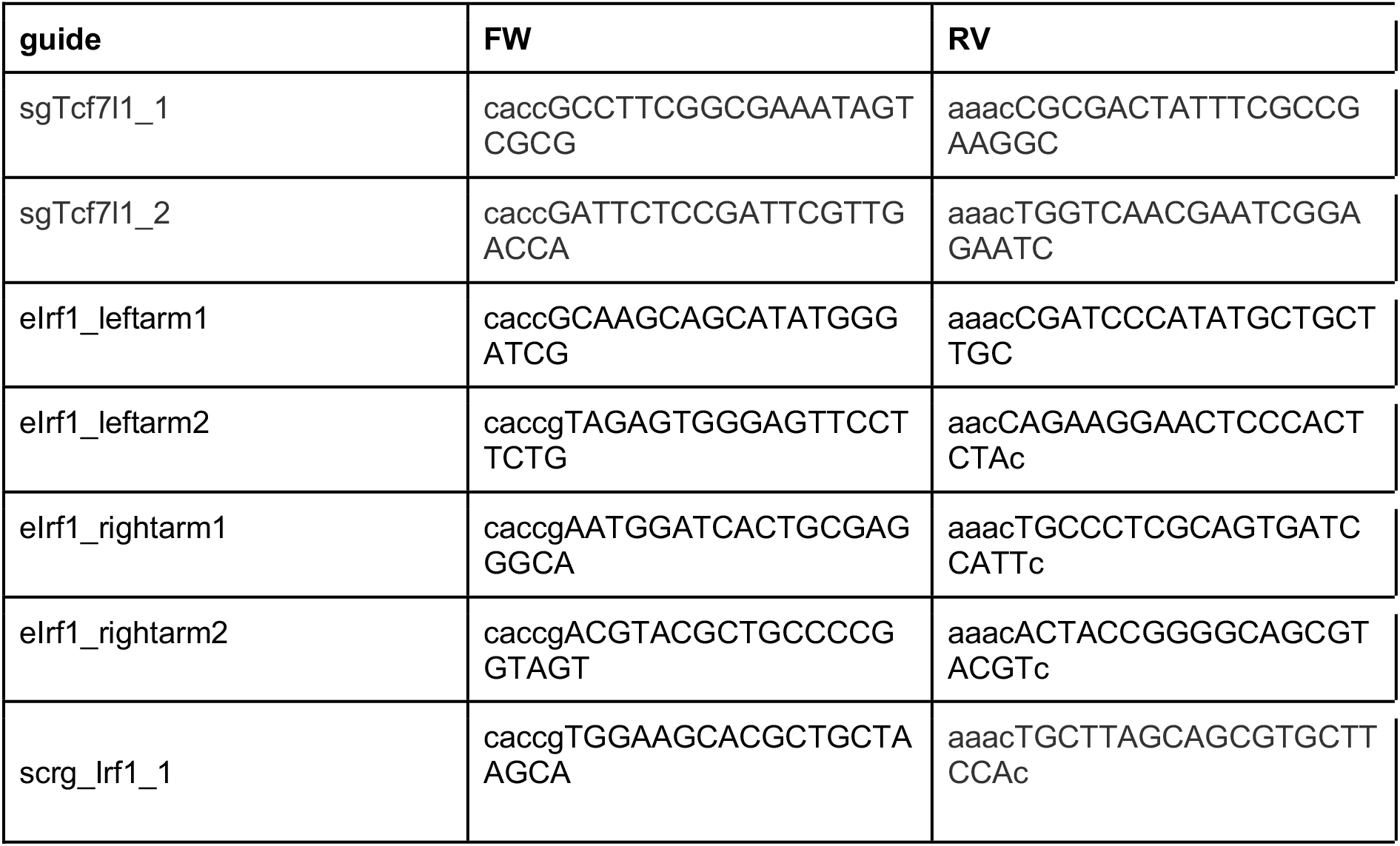
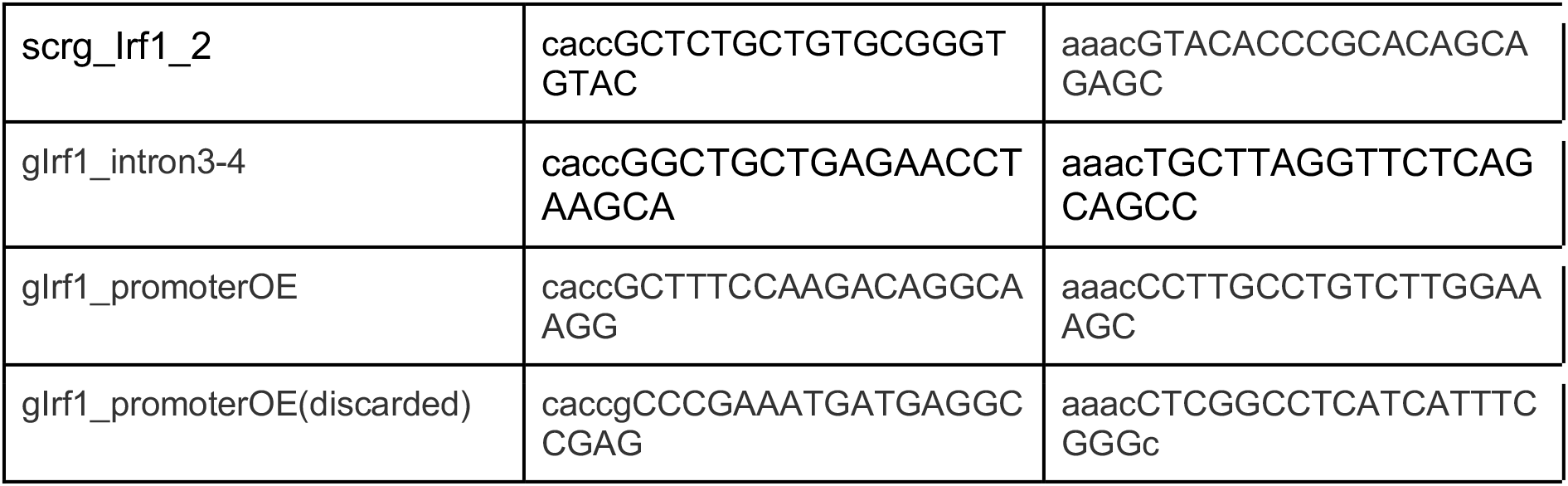

### antibodies

| name | source | usage |
| --- | --- | --- |
| GAPDH | Santa Cruz sc-365062 | WB 1:10000 |
| lrf1 | CST #8478 | IF 1:200, WB 1:1000 |
| Otx2 | R&D Systems AF1979 | WB 1:1000 |
| vinculin | Santa Cruz sc-73614 | WB 1:5000 |

### used datasets

|  |  |
| --- | --- |
| ChIP: H3K27ac, Oct4, p300 | GSE56138 |
| lrf1 ChIP in BMDM | GSE77886 |
| human scRNA pre-implantation embryo | E-MTAB-3929 |
| mouse scRNA gastrulation time series | https://marionilab.cruk.cam.ac.uk/MouseGastrulation2018/ |

All NGS data generated in this study will be made available.

---

**Figure S1 related to Figure 1.**
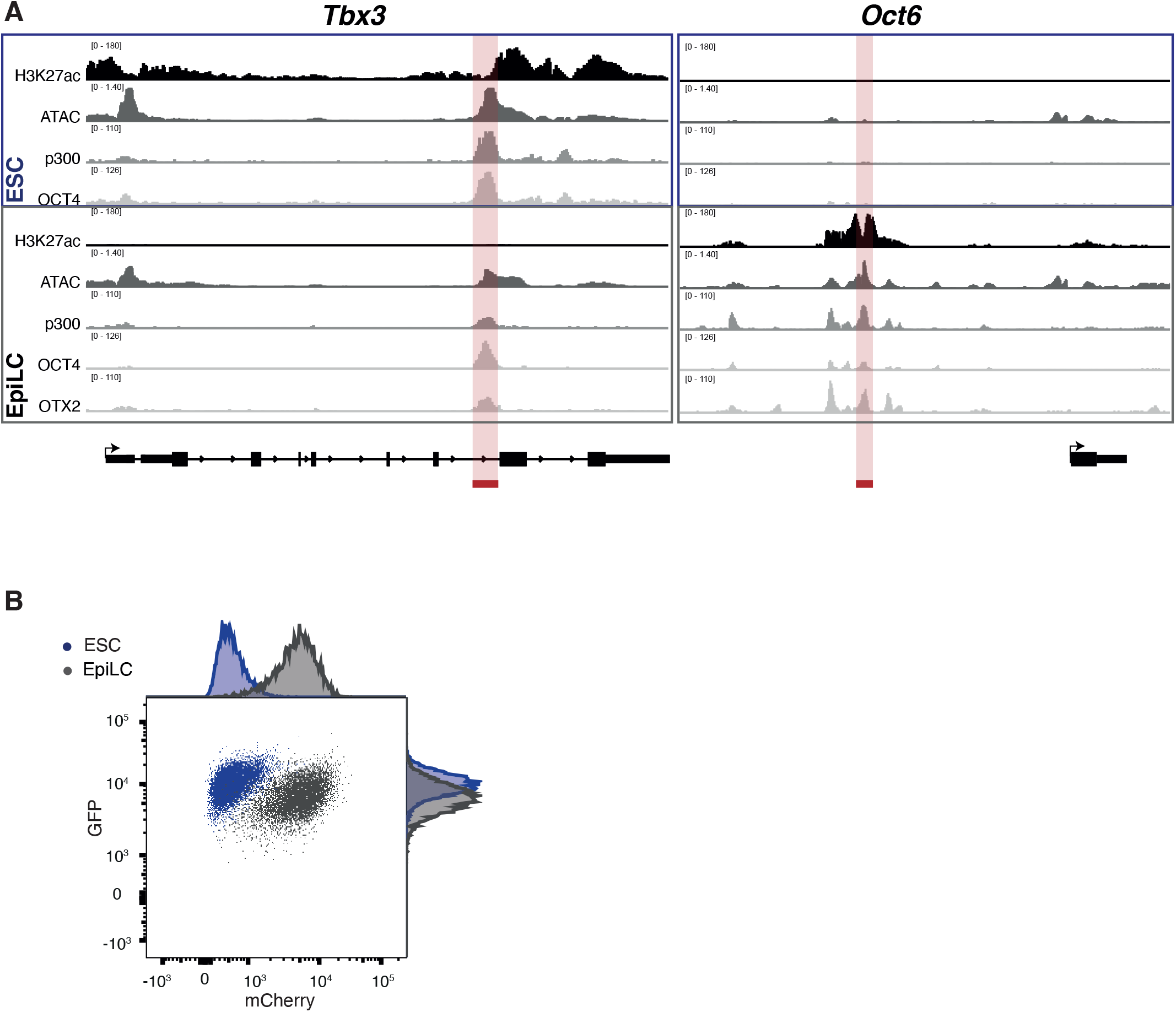
**A** Chromatin context of *Tbx3* and *Oct6* loci in ESC and EpiLC. Enhancer regions used as reporters are indicated by red boxes. ChIP data was generated by Buecker et al. 2014. **B** Representative flow cytometry profiles of reporter cell lines in ESC and differentiated to EpiLC conditions, independent clone. See also Figure 1B.

**Figure S2 related to Figure 2.**
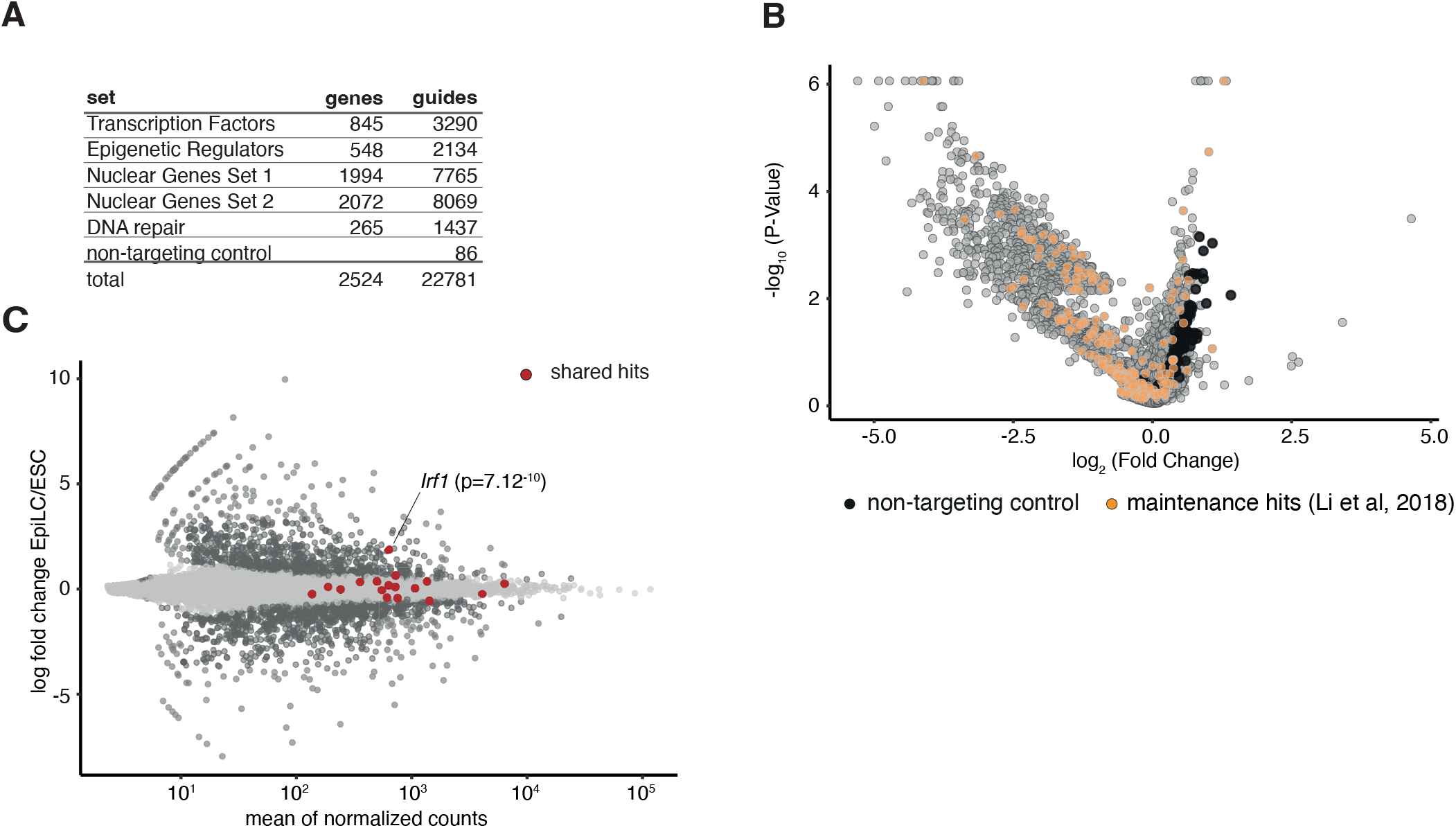
**A** Library information about genes and unique guides used in CRISPR-KO screen. **B** As in Figure 2B, guide representation in ESCs with CAS9 vs. ESCs without CAS9. Indicated are known ESC maintenance genes (Li et al. 2018). **C** MA plot of gene expression changes in EpiLC vs. ESC. The candidate factors which were identified as shared between both screening conditions (Figure 2E) are indicated in red. p-value for *Irf1* calculated with DESeq2.

**Figure S3.1 related to Figure 3.**
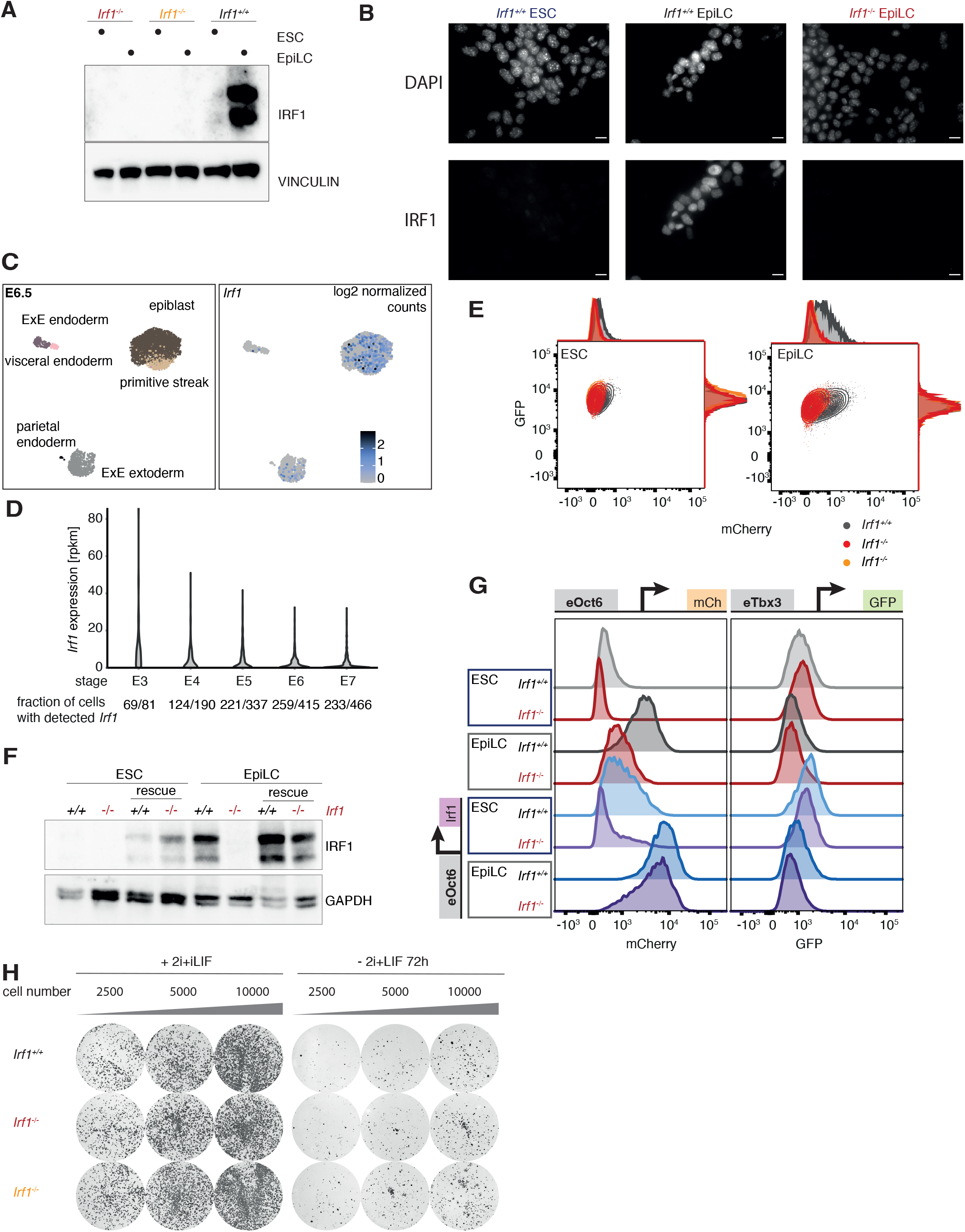
**A** Western Blot Analysis of *Irf1* KO in ESC and EpiLC, probed with antibodies against IRF1 and VINCULIN as loading control. Shown are two independent clones. **B** Immunofluorescence with IRF1 antibody of *Irf1^+/+^* and *Irf1^-/-^* ESC and EpiLC. Nuclei are stained with DAPI. Scale bars are 10 µm. **C** Cellular identity (left) and *Irf1* mRNA expression (right) in scRNA-seq data mouse embryo E6.5, data from (Pijuan-Sala et al. 2019) **D** *Irf1* mRNA expression in scRNA-seq data, human blastocyst of indicated stages. Fraction of cells in which *Irf1* reads were detected is indicated below. Data from Petropoulos et al. 2016. **E** Representative flow cytometry profiles of reporter cell lines in ESC and EpiLC conditions, *Irf1^+/+^* and *Irf1^-/-^*. Shown are two independent cell lines. **F** Western Blot analysis of ectopic *Irf1* expression in *Irf1^+/+^* and *Irf1^-/-^* background, in ESC and EpiLC cells, probed with antibodies against IRF1 and GAPDH as loading control. **G** Representative flow cytometry profiles of reporter cell lines in ESC and EpiLC condition, *Irf1^+/+^* and *Irf1^-/-^* and ectopic *Irf1* expression. **H** Alkaline phosphatase staining of *Irf1^+/+^* and *Irf1^-/-^* in different cell seeding densities. Left panel: cells were kept in 2i+LIF medium to control for seeding density, right panel: cells were differentiated for 72h before being placed back in 2i+LIF medium.

**Figure S3.2 related to Figure 3.**
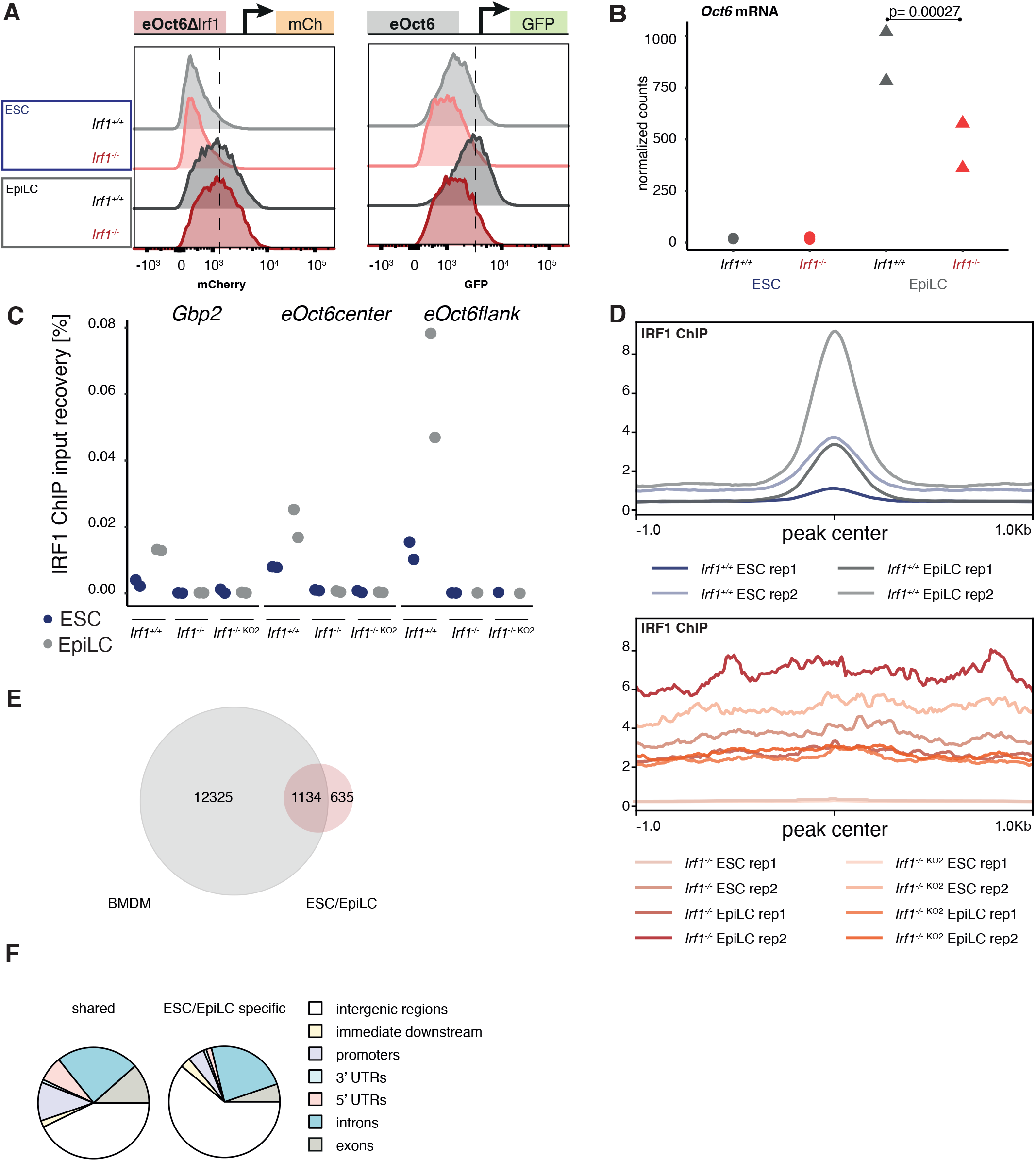
**A** As shown in Figure 3A: representative flow cytometry profiles of ESC and EpiLC cells, *Irf1^+/+^* and *Irf1^-/-^,* independent clone. *eOct6ΔIrf* and *eOct6* control mCherry and GFP, respectively. **B** Normalized counts for *Oct6* mRNA in *Irf1^+/+^* and *Irf1^-/-^* ESC and EpiLC Quant-Seq, two replicates were analyzed. p-value calculated by DESeq2. **C** ChIP-qPCR analysis for the *Gbp2* promoter as known IRF1 binding site and two primer sets for the *Oct6* enhancer. Values are calculated as input recovery [%]. **D** Signal strength around all identified IRF1 binding sites for indicated conditions. Top panel: *Irf1^+/+^* ESC and EpiLCs, bottom panel: two independent *Irf1^-/-^* cell lines in ESC and EpiLC conditions. **E** Venn diagram of overlap between chromatin IRF1 binding sites identified in bone marrow derived macrophages (BMDM) -INFɣ/+INFɣ (Langlais, Barreiro, and Gros 2016) and ESC/EpiLC. **F** Assigned chromatin regions of IRF1 binding sites shared between BMDMs and ESC/EpiLC (left) and IRF1 binding sites only detected in ESC/EpiLC (right).

**Figure S4 related to Figure 4.**
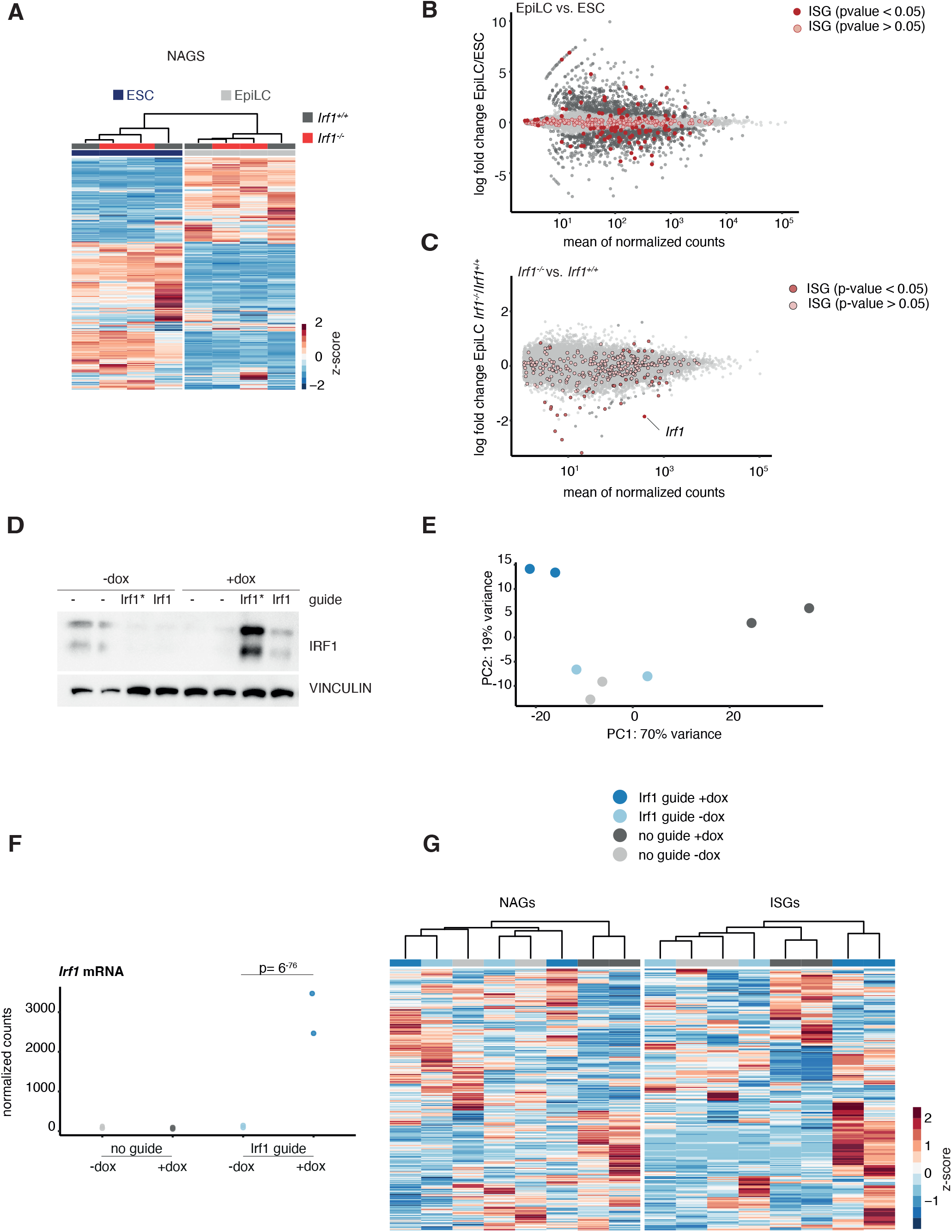
**A** Expression changes of naive associated genes (NAGs, Lackner et al. 2021) in *Irf1^+/+^* and *Irf1^-/-^* ESC and EpiLC based on QuantSeq RNA data. Data are shown as z-score. **B** MAplot of gene expression changes in EpiLC vs. ESC, ISGs are indicated. **C** MAplot of gene expression changes in EpiLC *Irf1^-/-^ vs. Irf1^+/+^,* ISGs are indicated. ISG which are differentially expressed (p value < 0.05) are plotted in Figure 4C. **D** Western Blot analysis of dox-inducible SunTag based IRF1 overexpression in ESCs, probed with antibodies against IRF1 and VINCULIN as loading control. Two different guides were tested, the guide marked with asterix (Irf1*) was used for RNA sequencing. **E** Principal component plot of dox-inducible SunTag-based IRF1 overexpression in ESCs, based on QuantSeq RNA data. **F** Normalized counts for *Irf1* mRNA in dox-inducible SunTag-based IRF1 overexpression in ESCs, based on QuantSeq RNA data. p-value calculated by DESeq2. **G** Expression changes of dox-inducible SunTag-based IRF1 overexpression, left panel NAGS, right panel full set of ISGs. Data are shown as z-score. Samples are color-coded as in Fig S4E.

**Figure S6 related to Figure 5.**
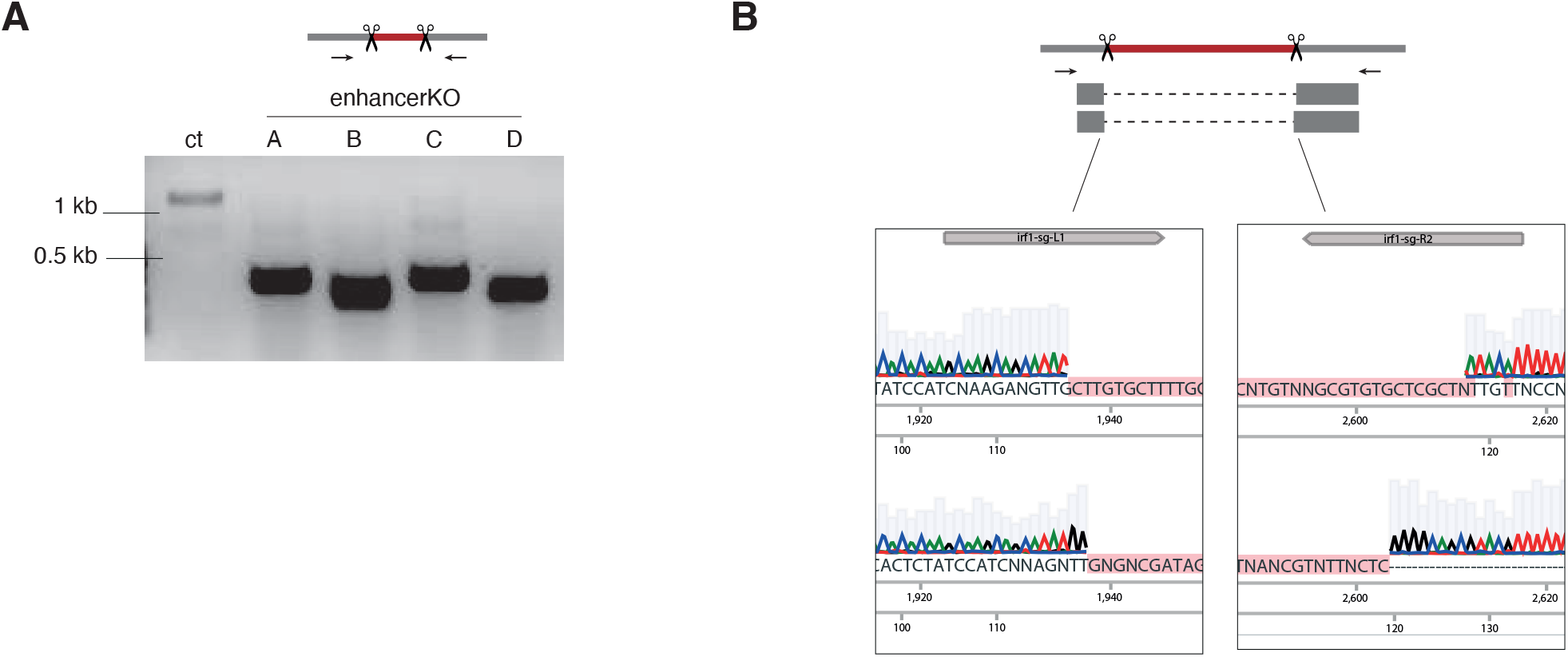
A Genotyping PCR of enhancer KO. Primers bind outside of editing site, producing smaller PCR products than on the control locus. Enhancer KO A and C were generated with one set of sgRNA, enhancer KO B and D with another set. B Examples for sanger sequencing of PCR products derived from Figure S6A, shown are enhancer KO B and D. Gapped alignment is indicated by dashed lines.

## Notes

### Competing Interest Statement

The authors have declared no competing interest.

## References

Bermingham, J R, S S Scherer, S O’Connell, E Arroyo, K A Kalla, F L Powell, and M G Rosenfeld. 1996. ‘Tst-1/Oct-6/SCIP Regulates a Unique Step in Peripheral Myelination and Is Required for Normal Respiration.’ Genes & Development 10 (14): 1751–62. https://doi.org/10.1101/gad.10.14.1751.

Betschinger, Joerg, Jennifer Nichols, Sabine Dietmann, Philip D. Corrin, Patrick J. Paddison, and Austin Smith. 2013. ‘Exit from Pluripotency Is Gated by Intracellular Redistribution of the BHLH Transcription Factor Tfe3’. Cell 153 (2): 335–47. https://doi.org/10.1016/j.cell.2013.03.012.

Buecker, Christa, Rajini Srinivasan, Zhixiang Wu, Eliezer Calo, Dario Acampora, Tiago Faial, Antonio Simeone, Minjia Tan, Tomasz Swigut, and Joanna Wysocka. 2014. ‘Reorganization of Enhancer Patterns in Transition from Naive to Primed Pluripotency’. Cell Stem Cell 14 (6): 838–53. https://doi.org/10.1016/j.stem.2014.04.003.

Burke, Derek C., Christopher F. Graham, and John M. Lehman. 1978. ‘Appearance of Interferon Inducibility and Sensitivity during Differentiation of Murine Teratocarcinoma Cells in Vitro’. Cell 13 (2): 243–48. https://doi.org/10.1016/0092-8674(78)90193-9.

Chen, Bohan, Chandan Gurung, Jason Guo, Chulan Kwon, and Yan-Lin Guo. 2020. ‘Pluripotent Stem Cells Are Insensitive to the Cytotoxicity of TNFα and IFNγ’. Reproduction 160 (4): 547–60. https://doi.org/10.1530/REP-20-0215.

Dunn, S.- J., G. Martello, B. Yordanov, S. Emmott, and A. G. Smith. 2014. ‘Defining an Essential Transcription Factor Program for Naive Pluripotency’. Science 344 (6188): 1156–60. https://doi.org/10.1126/science.1248882.

Festuccia, Nicola, Nick Owens, Almira Chervova, Agnès Dubois, and Pablo Navarro. 2021. ‘The Combined Action of Esrrb and Nr5a2 Is Essential for Murine Naïve Pluripotency’. Development 148 (17): dev199604. https://doi.org/10.1242/dev.199604.

Gao, X., P. Tate, P. Hu, R. Tjian, W. C. Skarnes, and Z. Wang. 2008. ‘ES Cell Pluripotency and Germ-Layer Formation Require the SWI/SNF Chromatin Remodeling Component BAF250a’. Proceedings of the National Academy of Sciences 105 (18): 6656–61. https://doi.org/10.1073/pnas.0801802105.

Grow, Edward J., Ryan A. Flynn, Shawn L. Chavez, Nicholas L. Bayless, Mark Wossidlo, Daniel J. Wesche, Lance Martin, et al. 2015. ‘Intrinsic Retroviral Reactivation in Human Preimplantation Embryos and Pluripotent Cells’. Nature 522 (7555): 221–25. https://doi.org/10.1038/nature14308.

Hackett, Jamie A., Yun Huang, Ufuk Günesdogan, Kristjan A. Gretarsson, Toshihiro Kobayashi, and M. Azim Surani. 2018. ‘Tracing the Transitions from Pluripotency to Germ Cell Fate with CRISPR Screening’. Nature Communications 9 (1): 4292. https://doi.org/10.1038/s41467-018-06230-0.

Harada, Hisashi, Keith Willison, Jun Sakakibara, Masaaki Miyamoto, Takashi Fujita, and Tadatsugu Taniguchi. 1990. ‘Absence of the Type I IFN System in EC Cells: Transcriptional Activator (IRF-1) and Repressor (IRF-2) Genes Are Developmentally Regulated’. Cell 63 (2): 303–12. https://doi.org/10.1016/0092-8674(90)90163-9.

Hart, Traver, Amy Hin Yan Tong, Katie Chan, Jolanda Van Leeuwen, Ashwin Seetharaman, Michael Aregger, Megha Chandrashekhar, et al. 2017. ‘Evaluation and Design of Genome-Wide CRISPR/SpCas9 Knockout Screens’. G3 Genes|Genomes|Genetics 7 (8): 2719–27. https://doi.org/10.1534/g3.117.041277.

Hayashi, Katsuhiko, Hiroshi Ohta, Kazuki Kurimoto, Shinya Aramaki, and Mitinori Saitou. 2011. ‘Reconstitution of the Mouse Germ Cell Specification Pathway in Culture by Pluripotent Stem Cells’. Cell 146 (4): 519–32. https://doi.org/10.1016/j.cell.2011.06.052.

Heurtier, Victor, Nick Owens, Inma Gonzalez, Florian Mueller, Caroline Proux, Damien Mornico, Philippe Clerc, Agnes Dubois, and Pablo Navarro. 2019. ‘The Molecular Logic of Nanog-Induced Self-Renewal in Mouse Embryonic Stem Cells’. Nature Communications 10 (1): 1109. https://doi.org/10.1038/s41467-019-09041-z.

Hofmann, Elisabeth, Ursula Reichart, Christian Gausterer, Christian Guelly, Dies Meijer, Mathias Müller, and Birgit Strobl. 2010. ‘Octamer-Binding Factor 6 (Oct-6/Pou3f1) Is Induced by Interferon and Contributes to DsRNA-Mediated Transcriptional Responses’. BMC Cell Biology 11 (1): 61. https://doi.org/10.1186/1471-2121-11-61.

Hsu, Patrick D, David A Scott, Joshua A Weinstein, F Ann Ran, Silvana Konermann, Vineeta Agarwala, Yinqing Li, et al. 2013. ‘DNA Targeting Specificity of RNA-Guided Cas9 Nucleases’. Nature Biotechnology 31 (9): 827–32. https://doi.org/10.1038/nbt.2647.

Kalkan, Tüzer, Susanne Bornelöv, Carla Mulas, Evangelia Diamanti, Tim Lohoff, Meryem Ralser, Sjors Middelkamp, Patrick Lombard, Jennifer Nichols, and Austin Smith. 2019. ‘Complementary Activity of ETV5, RBPJ, and TCF3 Drives Formative Transition from Naive Pluripotency’. Cell Stem Cell 24 (5): 785–801.e7. https://doi.org/10.1016/j.stem.2019.03.017.

Kell, Alison M., and Michael Gale. 2015. ‘RIG-I in RNA Virus Recognition’. Virology 479–480 (May): 110–21. https://doi.org/10.1016/j.virol.2015.02.017.

Kim, Kee-Pyo, Dong Wook Han, Johnny Kim, and Hans R. Schöler. 2021. ‘Biological Importance of OCT Transcription Factors in Reprogramming and Development’. Experimental & Molecular Medicine 53 (6): 1018–28. https://doi.org/10.1038/s12276-021-00637-4.

Kim, Kee-Pyo, You Wu, Juyong Yoon, Kenjiro Adachi, Guangming Wu, Sergiy Velychko, Caitlin M. MacCarthy, et al. 2020. ‘Reprogramming Competence of OCT Factors Is Determined by Transactivation Domains’. Science Advances 6 (36): eaaz7364. https://doi.org/10.1126/sciadv.aaz7364.

Lackner, Andreas, Robert Sehlke, Marius Garmhausen, Giuliano Giuseppe Stirparo, Michelle Huth, Fabian Titz-Teixeira, Petra Lelij, et al. 2021. ‘Cooperative Genetic Networks Drive Embryonic Stem Cell Transition from Naïve to Formative Pluripotency’. The EMBO Journal, March. https://doi.org/10.15252/embj.2020105776.

Langlais, David, Luis B. Barreiro, and Philippe Gros. 2016. ‘The Macrophage IRF8/IRF1 Regulome Is Required for Protection against Infections and Is Associated with Chronic Inflammation’. Journal of Experimental Medicine 213 (4): 585–603. https://doi.org/10.1084/jem.20151764.

Lara-Astiaso, David, Assaf Weiner, Erika Lorenzo-Vivas, Irina Zaretsky, Diego Adhemar Jaitin, Eyal David, Hadas Keren-Shaul, et al. 2014. ‘Chromatin State Dynamics during Blood Formation’. Science 345 (6199): 943–49. https://doi.org/10.1126/science.1256271.

Leeb, Martin, Sabine Dietmann, Maike Paramor, Hitoshi Niwa, and Austin Smith. 2014. ‘Genetic Exploration of the Exit from Self-Renewal Using Haploid Embryonic Stem Cells’. Cell Stem Cell 14 (3): 385–93. https://doi.org/10.1016/j.stem.2013.12.008.

Li, Meng, Jason S.L. Yu, Katarzyna Tilgner, Swee Hoe Ong, Hiroko Koike-Yusa, and Kosuke Yusa. 2018. ‘Genome-Wide CRISPR-KO Screen Uncovers MTORC1-Mediated Gsk3 Regulation in Naive Pluripotency Maintenance and Dissolution’. Cell Reports 24 (2): 489–502. https://doi.org/10.1016/j.celrep.2018.06.027.

Matsuda, Kazunari, Tomoyuki Mikami, Shinya Oki, Hideaki Iida, Munazah Andrabi, Jeremy M. Boss, Katsushi Yamaguchi, Shuji Shigenobu, and Hisato Kondoh. 2017. ‘ChIP-Seq Analysis of Genomic Binding Regions of Five Major Transcription Factors Highlights a Central Role for ZIC2 in the Mouse Epiblast Stem Cell Gene Regulatory Network’. Development 144 (11): 1948–58. https://doi.org/10.1242/dev.143479.

McLean, Cory Y, Dave Bristor, Michael Hiller, Shoa L Clarke, Bruce T Schaar, Craig B Lowe, Aaron M Wenger, and Gill Bejerano. 2010. ‘GREAT Improves Functional Interpretation of Cis-Regulatory Regions’. Nature Biotechnology 28 (5): 495–501. https://doi.org/10.1038/nbt.1630.

Michlits, Georg, Maria Hubmann, Szu-Hsien Wu, Gintautas Vainorius, Elena Budusan, Sergei Zhuk, Thomas R Burkard, et al. 2017. ‘CRISPR-UMI: Single-Cell Lineage Tracing of Pooled CRISPR–Cas9 Screens’. Nature Methods 14 (12): 1191–97. https://doi.org/10.1038/nmeth.4466.

Molotkov, Andrei, Pierre Mazot, J. Richard Brewer, Ryan M. Cinalli, and Philippe Soriano. 2017. ‘Distinct Requirements for FGFR1 and FGFR2 in Primitive Endoderm Development and Exit from Pluripotency’. Developmental Cell 41 (5): 511–526.e4. https://doi.org/10.1016/j.devcel.2017.05.004.

Moussaieff, Arieh, Matthieu Rouleau, Daniel Kitsberg, Merav Cohen, Gahl Levy, Dinorah Barasch, Alina Nemirovski, et al. 2015. ‘Glycolysis-Mediated Changes in Acetyl-CoA and Histone Acetylation Control the Early Differentiation of Embryonic Stem Cells’. Cell Metabolism 21 (3): 392–402. https://doi.org/10.1016/j.cmet.2015.02.002.

Oshiumi, Hiroyuki, Moeko Miyashita, Masaaki Okamoto, Yuka Morioka, Masaru Okabe, Misako Matsumoto, and Tsukasa Seya. 2015. ‘DDX60 Is Involved in RIG-I-Dependent and Independent Antiviral Responses, and Its Function Is Attenuated by Virus-Induced EGFR Activation’. Cell Reports 11 (8): 1193–1207. https://doi.org/10.1016/j.celrep.2015.04.047.

Petropoulos, Sophie, Daniel Edsgärd, Björn Reinius, Qiaolin Deng, Sarita Pauliina Panula, Simone Codeluppi, Alvaro Plaza Reyes, Sten Linnarsson, Rickard Sandberg, and Fredrik Lanner. 2016. ‘Single-Cell RNA-Seq Reveals Lineage and X Chromosome Dynamics in Human Preimplantation Embryos’. Cell 165 (4): 1012–26. https://doi.org/10.1016/j.cell.2016.03.023.

Pijuan-Sala, Blanca, Jonathan A. Griffiths, Carolina Guibentif, Tom W. Hiscock, Wajid Jawaid, Fernando J. Calero-Nieto, Carla Mulas, et al. 2019. ‘A Single-Cell Molecular Map of Mouse Gastrulation and Early Organogenesis’. Nature 566 (7745): 490–95. https://doi.org/10.1038/s41586-019-0933-9.

Platanitis, Ekaterini, Duygu Demiroz, Anja Schneller, Katrin Fischer, Christophe Capelle, Markus Hartl, Thomas Gossenreiter, Mathias Müller, Maria Novatchkova, and Thomas Decker. 2019. ‘A Molecular Switch from STAT2-IRF9 to ISGF3 Underlies Interferon-Induced Gene Transcription’. Nature Communications 10 (1): 2921. https://doi.org/10.1038/s41467-019-10970-y.

Ramsauer, K., M. Farlik, G. Zupkovitz, C. Seiser, A. Kroger, H. Hauser, and T. Decker. 2007. ‘Distinct Modes of Action Applied by Transcription Factors STAT1 and IRF1 to Initiate Transcription of the IFN- -Inducible Gbp2 Gene’. Proceedings of the National Academy of Sciences 104 (8): 2849–54. https://doi.org/10.1073/pnas.0610944104.

Sabapathy, K. 1997. ‘Regulation of ES Cell Differentiation by Functional and Conformational Modulation of P53’. The EMBO Journal 16 (20): 6217–29. https://doi.org/10.1093/emboj/16.20.6217.

Sachs, Parysatis, Dong Ding, Philipp Bergmaier, Boris Lamp, Christina Schlagheck, Florian Finkernagel, Andrea Nist, Thorsten Stiewe, and Jacqueline E. Mermoud. 2019. ‘SMARCAD1 ATPase Activity Is Required to Silence Endogenous Retroviruses in Embryonic Stem Cells’. Nature Communications 10 (1): 1335. https://doi.org/10.1038/s41467-019-09078-0.

Seruggia, Davide, Martin Oti, Pratibha Tripathi, Matthew C. Canver, Lucy LeBlanc, Dafne C. Di Giammartino, Michael J. Bullen, et al. 2019. ‘TAF5L and TAF6L Maintain Self-Renewal of Embryonic Stem Cells via the MYC Regulatory Network’. Molecular Cell 74 (6): 1148–1163.e7. https://doi.org/10.1016/j.molcel.2019.03.025.

Shi, Bingbo, Dengfeng Gao, Liang Zhong, Minglei Zhi, Xiaogang Weng, Junjun Xu, Junhong Li, et al. 2020. ‘IRF-1 Expressed in the Inner Cell Mass of the Porcine Early Blastocyst Enhances the Pluripotency of Induced Pluripotent Stem Cells’. Stem Cell Research & Therapy 11 (1): 505. https://doi.org/10.1186/s13287-020-01983-2.

Smith, Austin. 2017. ‘Formative Pluripotency: The Executive Phase in a Developmental Continuum’. Development 144 (3): 365–73. https://doi.org/10.1242/dev.142679.

Swartzendruber, D. E., and J. M. Lehman. 1975. ‘Neoplastic Differentiation: Interaction of Simian Virus 40 and Polyoma Virus with Murine Teratocarcinoma Cells in Vitro’. Journal of Cellular Physiology 85 (2): 179–87. https://doi.org/10.1002/jcp.1040850204.

Thomas, Henry F., Elena Kotova, Swathi Jayaram, Axel Pilz, Merrit Romeike, Andreas Lackner, Thomas Penz, et al. 2021. ‘Temporal Dissection of an Enhancer Cluster Reveals Distinct Temporal and Functional Contributions of Individual Elements’. Molecular Cell 81 (5): 969–982.e13. https://doi.org/10.1016/j.molcel.2020.12.047.

Wolf, Daniel, and Stephen P. Goff. 2009. ‘Embryonic Stem Cells Use ZFP809 to Silence Retroviral DNAs’. Nature 458 (7242): 1201–4. https://doi.org/10.1038/nature07844.

Wray, Jason, Tüzer Kalkan, Sandra Gomez-Lopez, Dominik Eckardt, Andrew Cook, Rolf Kemler, and Austin Smith. 2011. ‘Inhibition of Glycogen Synthase Kinase-3 Alleviates Tcf3 Repression of the Pluripotency Network and Increases Embryonic Stem Cell Resistance to Differentiation’. Nature Cell Biology 13 (7): 838–45. https://doi.org/10.1038/ncb2267.

Wu, Xianfang, Viet Loan Dao Thi, Yumin Huang, Eva Billerbeck, Debjani Saha, Hans-Heinrich Hoffmann, Yaomei Wang, et al. 2018. ‘Intrinsic Immunity Shapes Viral Resistance of Stem Cells’. Cell 172 (3): 423–438.e25. https://doi.org/10.1016/j.cell.2017.11.018.

Yang, Shen-Hsi, Munazah Andrabi, Rebecca Biss, Syed Murtuza Baker, Mudassar Iqbal, and Andrew D. Sharrocks. 2019. ‘ZIC3 Controls the Transition from Naive to Primed Pluripotency’. Cell Reports 27 (11): 3215–3227.e6. https://doi.org/10.1016/j.celrep.2019.05.026.

Yang, Shen-Hsi, Tüzer Kalkan, Claire Morissroe, Hendrik Marks, Hendrik Stunnenberg, Austin Smith, and Andrew D. Sharrocks. 2014. ‘Otx2 and Oct4 Drive Early Enhancer Activation during Embryonic Stem Cell Transition from Naive Pluripotency’. Cell Reports 7 (6): 1968–81. https://doi.org/10.1016/j.celrep.2014.05.037.

